# Inhibition of FicD-mediated AMPylation and deAMPylation by Isoprenoid Diphosphates

**DOI:** 10.64898/2025.12.08.693056

**Authors:** Aubrie M. Blevins, Wei Peng, Lisa N. Kinch, Zihan Monshad, Andrea G. Paredes, Christina Volz, Jared Rutter, Amanda K. Casey, Kevin G. Hicks, Kim Orth

## Abstract

FicD regulates Unfolded Protein Response (UPR) through reversible AMPylation and deAMPylation of BiP, an HSP70 chaperone and master regulator of the UPR. FicD activity is regulated by ER-stress, catalyzing BiP AMPylation under low stress conditions to hold inactive chaperone in reserve. In stressed cells, FicD deAMPylates BiP, acutely increasing its active pool to assist in protein folding. Variants in UPR machinery, including those in the FicD gene, are linked to hereditary diseases. Despite the known role of FicD in UPR, in-vivo regulation of its activity remains elusive, and identifying metabolites that alter FicD activity could prove useful pharmaceutically. We applied an unbiased high-throughput screening platform, known as <u>M</u>ass spectrometry Integrated with equilibrium <u>D</u>ialysis for the discovery of <u>A</u>llostery <u>S</u>ystematically (MIDAS), to identify novel small molecule metabolites that might regulate FicD activity. MIDAS revealed interactions between FicD and two mavelonate pathway intermediates : geranyl-pyrophosphate and farnesyl-pyrophosphate. Biochemical characterization indicates that both potently inhibit FicD-mediated AMPylation and deAMPylation. The crystal structure of FicD bound to farnesyl-pyrophosphate demonstrates a competitive inhibition mechanism, with the pyrophosphate adopting the alpha and beta phosphate positions of ATP and the hydrocarbon chain filling the nucleoside pocket. FicD variants previously appeared as biochemically indistinguishable, yet lead to different human pathologies. We demonstrate farnesyl-pyrophosphate inhibits FicD^R374H^ and FicD^R374C^ variants implicated in causing hereditary spastic paraplegia, but not the FicD^R371S^ variant associated with neonatal diabetes. This study furthers our understanding of FicD inhibitors and distinguishes disease causing variants, providing insight into pharmacological targeting of UPR activity.

**Significance Statement:** FicD regulates UPR signaling in metazoans by fine-tuning BiP chaperone capacity. Therefore, targeting FicD activity may be a tractable method of altering UPR signaling for therapeutic benefit. We identify geranyl- and farnesyl-pyrophosphate as specific FicD inhibitors. Notably, these small molecules differentially inhibit disease-causing variants of FicD. A structure of farnesyl-pyrophosphate bound to the FicD active site helps explain the differential inhibition of pathogenic variants and provides insight into interactions that can be differentially exploited for modifying FicD activity. Their composition provides a novel chemical foundation for future drug development efforts targeting FicD activity.

## Introduction

The Endoplasmic Reticulum (ER) is a major hub of protein folding, assembly, modification, and transport. ER protein folding machinery is continuously challenged by dynamic fluctuations in ER protein load, and cells have evolved signaling mechanisms to sense and appropriately respond to stressors of this process (1–4). The Unfolded Protein Response (UPR) is an adaptive signaling pathway that both senses and responds to disruptions in ER protein homeostasis, or proteostasis (1, 4–7). When unfolded protein clients accumulate in the ER, UPR signaling reestablishes proteostasis through multiple mechanisms, including enhancing ER protein folding capacity and reducing the number of ER protein clients. The essential HSP70 chaperone BiP (Hspa5) serves as the master regulator of UPR activation through reversible association and inhibition of the UPR transducers IRE1, ATF6, and PERK (7–12). Maladaptive UPR activation is involved in a wide range of disease processes, such as cancer, neurodegenerative disease, and metabolic disease (3, 4, 6, 13–15). Significant efforts have focused on targeting the UPR and ER proteostasis for therapeutic development (3, 6, 14–16).

In metazoans, the ER transmembrane protein FicD acts as a rheostat for BiP chaperone activity via post-translational AMPylation and deAMPylation (17–20). FicD is a bifunctional enzyme, catalyzing both AMPylation and deAMPylation of BiP in an ER stress-dependent manner. During homeostasis or low ER Stress, FicD acts as an AMPylator, generating a pool of inactive BiP (17, 19, 21). FicD mediated AMPylation of the residue T518 in BiP’s substrate binding domain impairs J-protein stimulated ATP hydrolysis, locking BiP in its ATP-bound, chaperone-incompetent confirmation (22). ER stress induces a shift in FicD activity from AMPylation to deAMPylation of BiP, thereby restoring its chaperone function and acutely increasing the pool of active BiP (21, 23, 24). This ER stress-dependent activity in FicD has been demonstrated in many cell culture models and animal models where FicD is especially important in terminally differentiated cells that experience repeated physiological stress (12, 17, 19, 21).

The dual catalytic activity of FicD is mediated by a single Fido (<u>Fi</u>c, <u>Do</u>c, and AvrB) domain with a conserved α-helical fold (20, 25, 26). Although other substrates can be used by Fido domains, the majority of Fido domain-containing enzymes, including FicD, use ATP and Mg^2+^ to catalyze AMPylation using a conserved motif: HPFx(D/E)GNGR_1_xxR_2_ (20, 21, 25–27) **(Fig 1A)**. A second conserved motif (S/TxxxE(G/N)) comprises an autoinhibitory helix (*α-inh*) in the FicD enzyme. The *α-inh* glutamate sidechain (E234 in FicD) competes for the binding of the γ-phosphate of ATP with the sidechain of conserved arginine (R374 in FicD) in the catalytic motif **(Fig 1A, blue R, red E).** A salt bridge formed between E324 and R374 shifts FicD activity to deAMPylation (26, 28). Mutation of this glutamate to glycine (i.e. FicD^E234G^) breaks this salt bridge and abolishes this intramolecular autoinhibition, thereby, making FicD a constitutive AMPylator and abolishing deAMPylation activity (26, 29). The lack of regulation by *α-inh* is similar to the activity mediated by many bacterial pathogens that encode Fic-mediated activity, including VopS and AvrB (25, 30).

**Figure 1.**
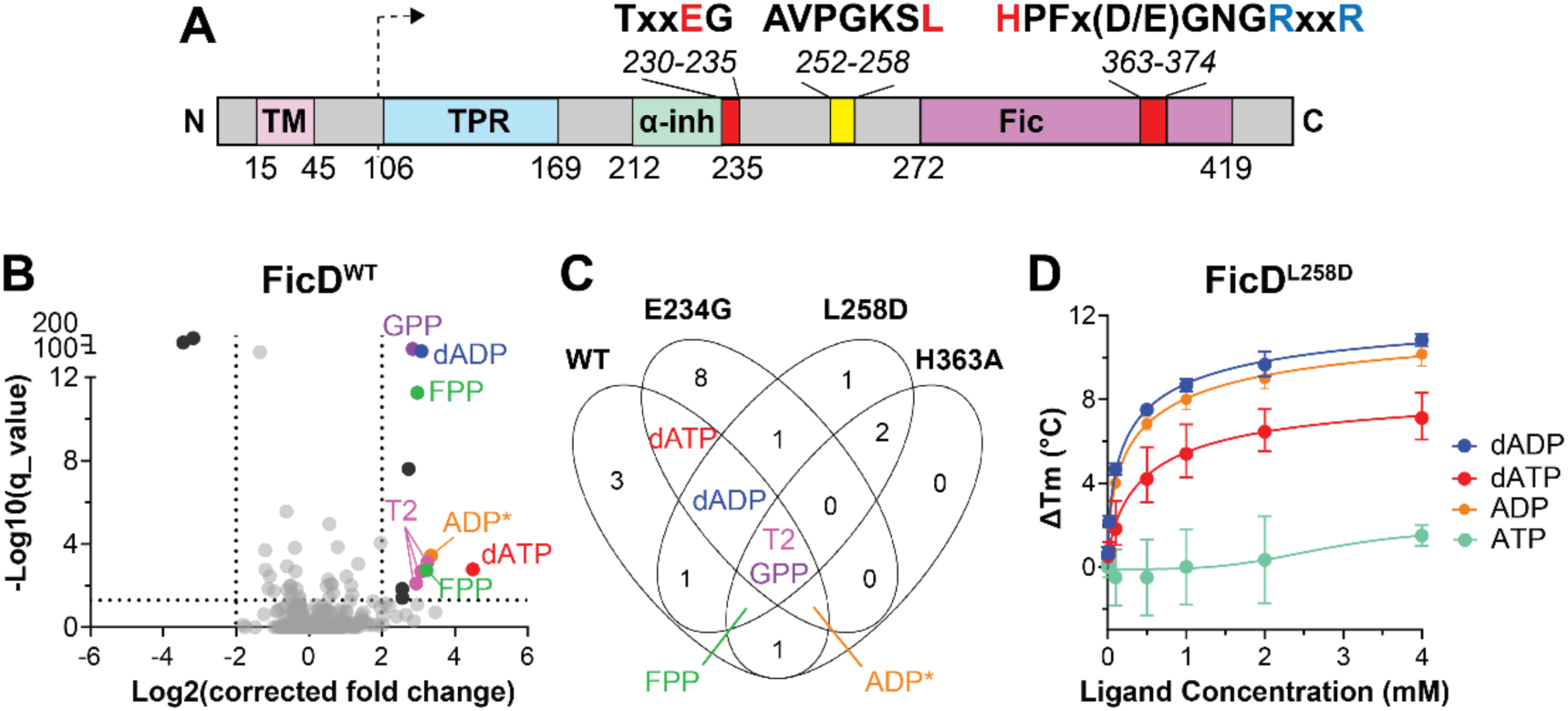
High-throughput FicD metabolite screen identified canonical ligands. **(A)** Domain diagram of FicD with labeled motifs that include functional (red) and pathogenic (blue) positions and the dotted arrow indicating the start site for expression constructs. **(B)** Volcano plot of FicD^WT^ MIDAS screen. Dotted lines indicate significance cutoffs (*Q* < 0.05 and |Log_2_ corrected fold change)| > 2). Significant PMIs are labeled and colored: GPP (purple), FPP (green), dADP (blue), dATP (red), ADP* for 3’,5’-ADP, ADP, or dGDP (orange), T2 (pink). Other PMIs are colored: significant but in <3 FicD conseruct screens (black) and nonsignificant (gray). **(C)** Venn diagram comparing significant PMIs detected in screens with WT, E234E, L258D, H363A. PMIs that overlap in 3 or more construct screens are labeled and colored as in B with nucleotides: ADP (orange), ATP (teal), dADP (blue), and dATP (red). Error bars represent three independent experiments.

The oligomeric state of the metazoan FicD enzyme also regulates the AMPylation/deAMPylation activity. The enzyme forms a dimer in solution, and mutations in the dimer interface (i.e. FicD^L258D^) yield a monomeric enzyme **(Fig 1A, red L).** The monomerization of FicD results in increased flexibility of *α-inh,* which facilitates both AMPylation competent binding of ATP and increases in AMPylation activity relative to wild-type FicD (24, 31). However, the FicD^L258D^ mutant, in contrast to the constitutive AMPylator FicD^E234G^, retains the ability to deAMPylate (24, 31). Collectively, these studies have revealed the structural relevance of the *α-inh* regulatory salt bridge and dimerization of FicD.

The importance of the inherent switching activity from a deAMPylator to an AMPylator is further realized when studying virulence factors and disease variants of metazoan FicD. For example, in some bacterial Fic enzymes, the *α-inh* and the Fic domain encompass two separate proteins and are expressed as a toxin-antitoxin module (27). AMPylation activity is regulated by the ratio of toxin to antitoxin. In a *Neisseria* Fic enzyme, autoAMPylation relieves this intramolecular autoinhibition, facilitating AMPylation activity (32). In *Enterococcus faecalis,* the Fic enzyme is regulated by levels of Mg^2+^ and Ca^2+^ to control its AMPylation/deAMPylation activity (33). In vitro, Ca^2+^ appears to inhibit the human FicD deAMPylation of BiP, but the physiological relevance remains unclear (33).

Studies in model organisms have illuminated both tissue and stress-dependent roles of FicD activity. In Drosophila, FicD (dFic) is required to maintain visual transmission in the eye (34). Flies deleted for dFic (dfic^-/-^ flies) sustain increased eye damage and an impaired ability to recover from repetitive light stress (21) and locomotor impairment (35). In FicD^-/-^ mice, a pharmaceutical induction of UPR stress caused increased fibrosis in the pancreas (21) and reduced ER stress adaptation in the liver (36). Conversely, enhanced UPR stress in the absence of FicD protected against hypertrophy-induced heart failure in mice (37). Additionally, *C. elegans* lacking Fic models (*fic-1^-/-^*) show enhanced UPR induction and chaperone expression that limited polyglutamine toxicity (38). These studies suggest that targeting FicD activity in specific tissues may provide a beneficial strategy by altering UPR signaling.

Missense mutations in the FicD active site are linked to pathologies in human patients **(Fig 1A, blue Rs)**. A rare homozygous recessive FicD^R371S^ variant is associated with neonatal diabetes and severe neurodevelopmental impairment (39). A preclinical mouse model of this disease indicates FicD^R371S^ dysregulation causes progressive loss of islet organization and impaired insulin secretion (40). Two additional rare homozygous FicD variants, FicD^R374C^ and FicD^R374H^, are implicated in hereditary spastic paraplegia and a risk of diabetes in adulthood (41, 42). Notably, although the recessive FicD variants have similar catalytic activity with weakened AMPylation and no deAMPylation activity, they cause different pathologies. Interestingly, in both flies and mice, the FicD^E234G^ allele that encodes strong, constitutive AMPylation activity is a homozygous lethal variant (24, 43). The accumulation of AMPylated BiP caused by a loss of FicD deAMPylation activity in FicD^R371S^, FicD^R374H^, and FicD^R374C^ patients is likely central to disease progression, which highlights the possible benefits of targeting pathologic FicD activity.

In this work, we utilize an unbiased, high-throughput screen known as <u>M</u>ass spectrometry Integrated with equilibrium <u>D</u>ialysis for the discovery of <u>A</u>llostery <u>S</u>ystematically (MIDAS), to investigate the possibility of small molecule metabolite regulation of FicD activity (44). Our screen identified the isoprenoid diphosphates geranyl-pyrophosphate (GPP) and farnesyl-pyrophosphate (FPP) as potent inhibitors of FicD-mediated AMPylation and deAMPylation. Orthogonal binding assays and in-vitro activity assays reveal that the interactions between GPP/FPP and FicD are specific to the metazoan enzymes, as GPP/FPP is unable to inhibit the activities of bacterial proteins with Fido domains. A crystal structure of metazoan FicD bound to FPP reveals that FPP binds within the FicD active site in a manner analogous to ADP binding. Finally, we generated recombinant proteins for the human FicD variants known to cause human disease and observed FPP binding and inhibition for FicD^R374H^ and FicD^R374C^ variants, but not for the FicD^R371S^ variant. Overall, our work demonstrates unprecedented regulation of the Fido domain by a novel class of small molecules, which may provide a scaffold for the development of specific FicD inhibitors.

## Results

### MIDAS screen reveals putative small-molecule metabolites that interact with FicD

To identify novel small-molecule metabolite interactions with FicD, we applied an unbiased high-throughput screening approach known as MIDAS (44). Given the high concentrations of protein required for MIDAS screening, we purified the soluble ER luminal domain (residues 105-458) of wild-type human FicD and three mutants FicD^E234G^, FicD^L258D^, and FicD^H363A^ for screening (**Fig 1A, dotted arrow**). These mutants were selected based on their well-established biochemical features (23, 24, 31). All three are predicted to enhance detection of protein-metabolite interactions (PMIs) relative to FicD^WT^ through increased affinity for nucleotide substrates (FicD^E234G^), improved solubility (FicD^L258D^), and loss of catalytic turnover (FicD^H363A^) (**Fig 1A)**. To decrease the potential for nonspecific small-molecule binding, we ensured the purity and homogeneity of our FicD preparations using SDS-PAGE of purified proteins (**Fig S1A and S1B**). As reported by other investigators (24, 31), we observe an increase in elution volume of the FicD^L258D^ mutant, consistent with monomerization (**Fig S1C**). Finally, intact mass of screened proteins did not reveal obvious contamination (**Fig S1D**).

Out of the 602 naturally occurring human metabolites in the MIDAS library, we applied a significance cutoff (q value <0.05) to identify 12 PMIs with a fold change of >2 and two PMIs with a fold change <-2 in the FicD^WT^ screen (**Fig 1A and Table S1**). Hits were additionally triaged based on detection of multiple adducts of the metabolite in a single dataset and identification of the same metabolites across three or more of the FicD construct screens (**Fig 1B**, Venn diagram overlap). Three chemical classes passed this initial triage: adenosine nucleotides, isoprenoid diphosphates, and thyroid hormone intermediates (**Fig 1A and 1B, Tables S1-S4**). These hits were all significantly enriched, indicating a stable binding that does not result in catalytic turnover.

Two of the chemical classes that passed initial triage represent known FicD PMIs: adenosine nucleotides and thyroid hormone intermediates. FicD uses ATP and Mg^2+^ to catalyze AMPylation. ADP readily binds to FicD, whereas intramolecular autoinhibition of AMPylation activity destabilizes ATP binding (26, 28, 39). In agreement with the canonical activity of FicD, an adduct consistent with ADP and the closely related analog, deoxyadenosine diphosphate (dADP), were significantly enriched in three of the four FicD constructs. The adduct consistent with ADP was identified in FicD^WT^, FicD^E234G^, and FicD^H363A^ screens, while dADP was identified in FicD^WT^, FicD^E234G^, and FicD^L258D^ screens (**Fig 1A and Tables S1-S4**). The FicD substrate ATP, although it did not pass the significance cutoff, was only detected in the screen with FicD^E234G^, which is not intrinsically autoinhibited, (**Dataset 2**). However, the dATP analog was significantly enriched in the FicD^WT^ and FicD^E234G^ datasets (**Fig 1A and Tables S1-S2**). dATP was also enriched in the FicD^L258D^ dataset, although it did not pass the significance cutoff (**Dataset 3**). Finally, previous drug screens of FicD have reported liothyronine, a thyroid hormone intermediate, as a small-molecule inhibitor of FicD (45). Consistent with these findings, 3,3’-Diiodothyronine was significantly enriched in all FicD mutant datasets. Overall, the detection of canonical FicD ligands served as internal positive controls for MIDAS screening, which supported our confidence in the previously unreported interactions discussed below.

A few spurious hits were enriched or depleted in only one FicD construct dataset (**Fig 1A, Tables S1-S4**). For example, the small, acidic compound glyoxylic acid was significantly depleted in FicD^WT^ (**Table S1**), but not in the FicD mutant datasets (**Table S2-S4**). In MIDAS, depletion may indicate either a catalytic turnover that is faster than the rate of diffusion across the dialysis membrane, a non-covalent interaction that resists release from the interacting protein during methanol precipitation, or a covalent modification of the protein. Accordingly, glyoxylic acid is known to spontaneously and non-enzymatically react with protein sidechains, a modification known as glycation (46).

### MIDAS identifies analogues of canonical FicD ligands

MIDAS screening identified putative interactions between FicD and dADP/dATP, which are the deoxy form of canonical FicD ligands ADP and ATP. To validate the interaction between FicD and dADP/dATP, we performed differential scanning fluorimetry (DSF) to measure alterations in FicD’s thermal stability upon ligand binding. The monomeric FicD^L258D^ mutant produced a prototypical sigmoidal melt curve reflecting the melting temperature (T_m_) of a single unfolding transition state. Consistent with the MIDAS screen, addition of increasing dADP or dATP ligand concentrations lead to higher T_m_ values, indicating stabilization of the protein and binding of the metabolites (**Fig S2A-B**).

The dose response curves for FicD^L258D^ in the presence of dADP and dATP reached maximal shifts (∼13 °C and ∼9 °C, respectively) at saturating ligand concentrations, with EC50s (0.28 mM and 0.53 mM, respectively) (**Fig 1D**). The dose response curve for dADP mirrored that of ADP (∼13°C shift with EC50 0.37 mM, **Fig 1D**, orange curve), indicating a similar binding mode for the two ligands. Consistent with previous reports (26, 31), the binding curve for ATP did not show saturation at the indicated concentrations (**Fig 1D**, green curve). The weak binding of ATP to FicD^L258D^ is consistent with previous observations that *α-inh* flexibility conferred by the L258D mutation alters the ATP binding mode without altering ATP affinity (31). In contrast to ATP, the stable binding of dATP to FicD^L258D^ suggests *α-inh* helix can accommodate the ligand with an increased affinity.

Because monomerization-induced flexibility of the FicD^L258D^ *α-inh* helix increases its ability to accommodate nucleotide ligands (26, 31), we used DSF to verify nucleotide binding to FicD^WT^. While the monomeric FicD^L258D^ displayed a monophasic melt transition with a T_m_ of 49.5°C (**Fig S2A and S2B**), the melt transition for dimeric FicD^WT^ was more complex, with a ∼12°C shift toward higher temperature (**Fig S2C and S2D**). This T_m_ difference is consistent with previous reports comparing the monomeric and dimeric enzymes (31). The dose response of ADP, dADP, and dATP for the FicD^WT^ (**Fig S2A-S2E**) follow a similar trend to those for FicD^L258D^, with the loss of the 2’ hydroxyl trending towards slight improvements in binding. The minor differences in nucleotide affinity are consistent with previous reports, which note that monomerization does not alter ATP binding affinity, but rather it involves the positioning of ATP in the FicD active site (31). Given that FicD^L258D^ retains bifunctional AMPylation and deAMPylation activities with only modest differences in nucleotide binding, we leveraged the monophasic melt transition of the L258D mutant to facilitate dose-response analysis.

### Geranyl-pyrophosphate and Farnesyl-pyrophosphate are specific FicD inhibitors

MIDAS screening identified the isoprenoid diphosphate GPP to be significantly enriched in all four FicD mutant datasets, and the related metabolite FPP was significantly enriched in FicD^WT^, FicD^L258D^, and FicD^H363A^ **(Fig 1A, Tables S1-S4)**. Further support for these PMIs is reflected by multiple adducts of GPP and FPP being significantly enriched. GPP and FPP are pyrophosphate esters of the terpenoids geraniol and farnesol (47). GPP contains two isoprene units and FPP contains three isoprene units (**Fig 2A**). In eukaryotes, they are intermediates of the mevalonate pathway, which produces isoprenoids, sterols, and lipids **(Fig. 2A)** (47, 48). In DSF, GPP and FPP elicited a dose-dependent and saturating increase in the τιT_m_ of FicD^L258D^, yielding EC50s of 14.8uM and 9.6uM, respectively (**Fig 2B**). Notably, these apparent binding constants are ∼20-30 fold tighter than that of ADP.

**Figure 2.**
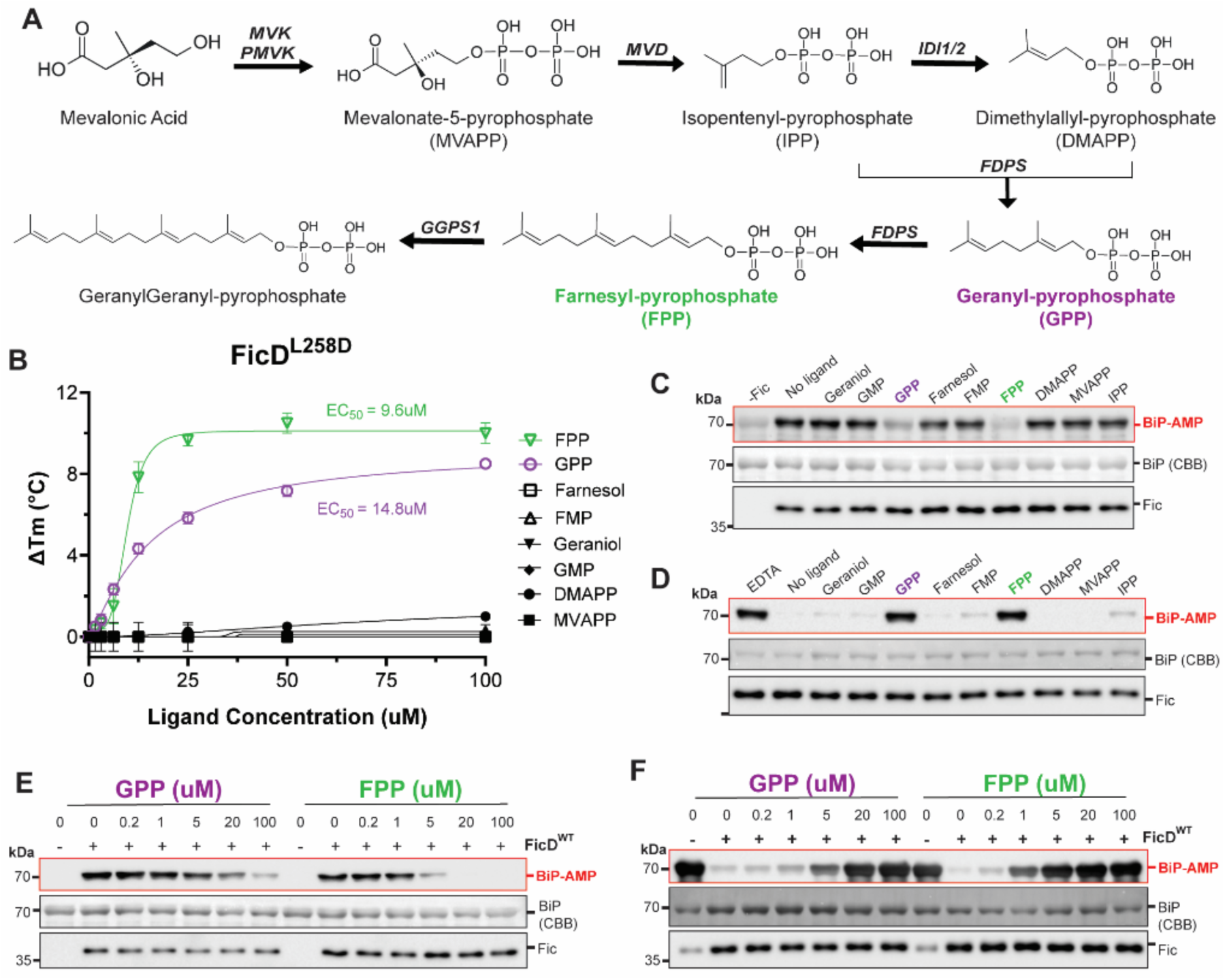
Geranyl-pyrophosphate and Farnesyl-pyrophosphate are specific FicD inhibitors. **(A)** Mevalonate Pathway for GPP and FPP biosynthesis in humans. Structures were created in MarvinSketch. **(B)** DSF melting of FicD^L258D^ with the indicated metabolites (legend) in the presence of 5mM MgCl_2_. Error bars represent two or three independent experiments. **(C and D)** Western blot of **(C)** Δ26hBiP^T229A^ AMPylation and **(D)** deAMPylation by FicD^WT^ in the presence of metabolites (100uM) or EDTA (5mM). **(E and F)** Western blots of GPP (left) or FPP (right) dose response inhibition of **(E)** Δ26hBiP^T229A^ AMPylation and **(F)** deAMPylation by FicD^WT^. AMPylated BiP and FicD were quantified by immunodetection and total BiP was quantified by CBB in all activity assays.

To ensure specificity of binding to the FicD enzyme, we performed DSF with Glutathione S-transferase (GST) and BiP in the presence of the same increasing metabolite concentrations. Neither GPP nor FPP elicited a change in the GST melt curves or T_m_ (**Fig S3**) at the same concentrations measured for FicD. Like FicD, BiP is an ATP/Mg^2+^ dependent protein that could potentially leverage the same binding mode as GPP/FPP. However, neither GPP nor FPP elicited a change the melt curves or T_m_ of BiP (**Fig S4**). Together, these results indicate that binding of GPP/FPP at the evaluated concentrations is specific for the FicD protein.

To probe the structural specificity of the GPP/FPP metabolites, we tested binding of various analogs to FicD^L258D^ using DSF. The farnesol and geraniol isoprenoid chains lacking pyrophosphate did not bind to FicD^L258D^, nor did the mono-phosphate versions: geranyl-monophosphate (GMP) and farnesyl-monophosphate (FMP) (**Fig 2B**). These results demonstrate the importance of the diphosphate in conferring binding to FicD. Similarly, dimethylallyl-pyrophosphate (DMAPP) composed of a single isoprene unit with a pyrophosphate did not bind to FicD^L258D^. Moreover, mevalonate-pyrophosphate (MVAPP), which is the precursor to DMAPP, also did not bind to FicD (**Fig 2B**). Together, these binding studies indicate that the specific combination of at least two isoprene units and a diphosphate are required for FicD binding.

We next sought to assess the effects of FPP, GPP, and isoprenoid analogues on FicD activity. While GPP and FPP at 100uM concentration inhibited the AMPylation activity of FicD^WT^, the analogs geraniol, farnesol, GMP, FMP, DMAPP, and MVAPP did not affect FicD^WT^ AMPylation activity at the same concentrations (**Fig 2C**). Similarly, GPP and FPP at 100uM concentration inhibited the deAMPylation activity of FicD^WT^ (**Fig 2D**). However, geraniol, farnesol, GMP, FMP, DMAPP, and MVAPP did not affect FicD^WT^ deAMPylation activity at the same concentrations (**Fig 2D**). These results corroborate the findings of our DSF experiments (**Fig 2B**).

We then assayed the potency of GPP/FPP induced inhibition of AMPylation and deAMPylation activity. In-vitro AMPylation and deAMPylation assays revealed dose-dependent inhibition of both AMPylation and deAMPylation activities of FicD (**Fig 2E and 2F**). Consistent with our DSF data, FPP was a more potent inhibitor of both AMPylation and deAMPylation activity than GPP. FPP inhibition of AMPylation and deAMPylation started as low as 5uM and 1uM, respectively (**Fig 2E and 2F**, right panels), while GPP inhibition of AMPylation and deAMPylation activity started as low as 20uM and 5uM, respectively (**Fig 2E and 2F**, left panels).

Finally, we explored the inhibitory effects of GPP/FPP on other Fido proteins to see if these metabolites could serve as generic inhibitors of the Fido domain. To test this specificity, we performed in-vitro activity assays with GPP/FPP and two constitutively active Fido proteins from bacteria: VopS from *Vibrio parahaemolyticus* (25) and AvrB from *Pseudomonas syringae* (30). Neither GPP nor FPP affected the AMPylation activity of VopS (**Fig S5A**). Similarly, neither GPP nor FPP affected the rhamnosylation activity of AvrB (**Fig S5B**).

DSF measurements suggested high-affinity binding of FPP and GPP to FicD when compared to nucleotides (**Fig 1B and 2B**). To further support this notion, we used ITC to determine the precise K_d_ of FPP binding. We observed FicD^L258D^ bound to FPP with a K_d_ of approximately 60 nM at a MgCl_2_ concentration of 4mM (**Fig 3A**). In these experiments, the interaction between FPP and Mg^2+^ generated endothermic peaks that interfered with detecting the interaction between FPP, Mg^2+^, and FicD (**Fig 3A**, upper panels). The stoichiometry of FPP binding to FicD was consistently <0.85, which may indicate that Mg^2+^ was not saturating. Because the endothermic interaction between Mg^2+^ and FPP limited the quantity of MgCl_2_ we could add, the measured K_d_ could be larger than the true value. Nevertheless, the K_d_ of FPP binding was observed to be over one order of magnitude smaller than that for ADP, which is reported to be 1.5uM for FicD^WT^ (26).

**Figure 3.**
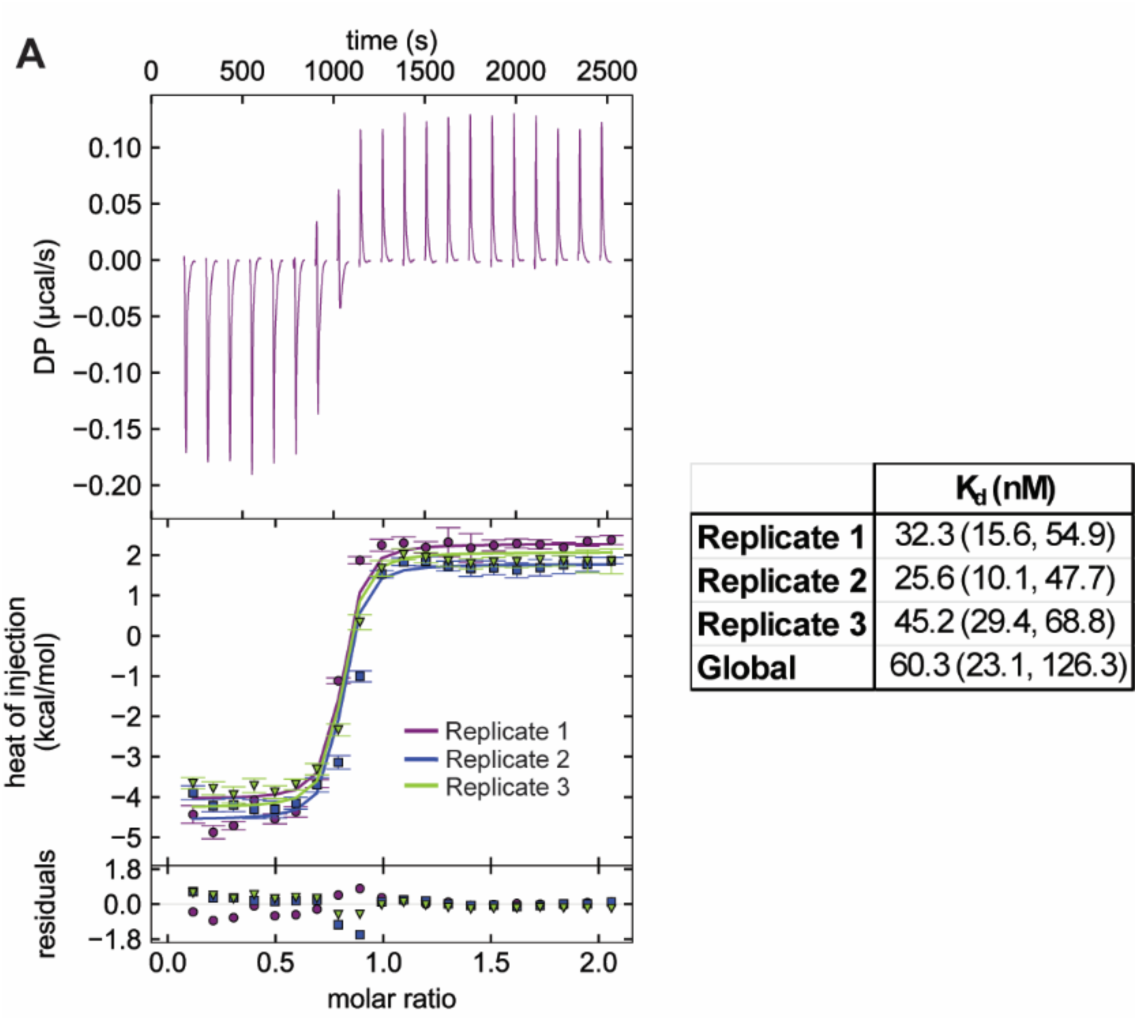
ITC with FicD and Farnesyl-pyrophosphate. **(A)** Representative ITC thermogram for FPP binding to Δ104hFicD^L258D^ (upper panel). Heats of injection of all three replicates are displayed (lower panel). Dissociation constants (K_d_) of individual replicates and global fitting of triplicate isotherms are reported with 1σ error intervals in parenthesis.

### Isoprenoid -pyrophosphate binding to FicD requires Mg^2+^

Our biochemical data indicates that FPP is a potent and specific inhibitor of FicD, whose binding and inhibition requires the pyrophosphate moiety. Given this requirement, we considered if the inhibitor pyrophosphate bound in the active site of the enzyme in a manner analogous to that of the nucleotide, which coordinates the α- and β- phosphates together with Mg^2+^ (26). Accordingly in DSF, the ADP, dADP, ATP, and dATP nucleotides did not bind to FicD^L258D^ in the absence of MgCl_2_ (**Fig 4A**). Conversely, the addition of 5mM MgCl_2_ conferred ADP, dADP, and dATP binding to FicD^L258D^ (**Fig 4B**).

**Figure 4.**
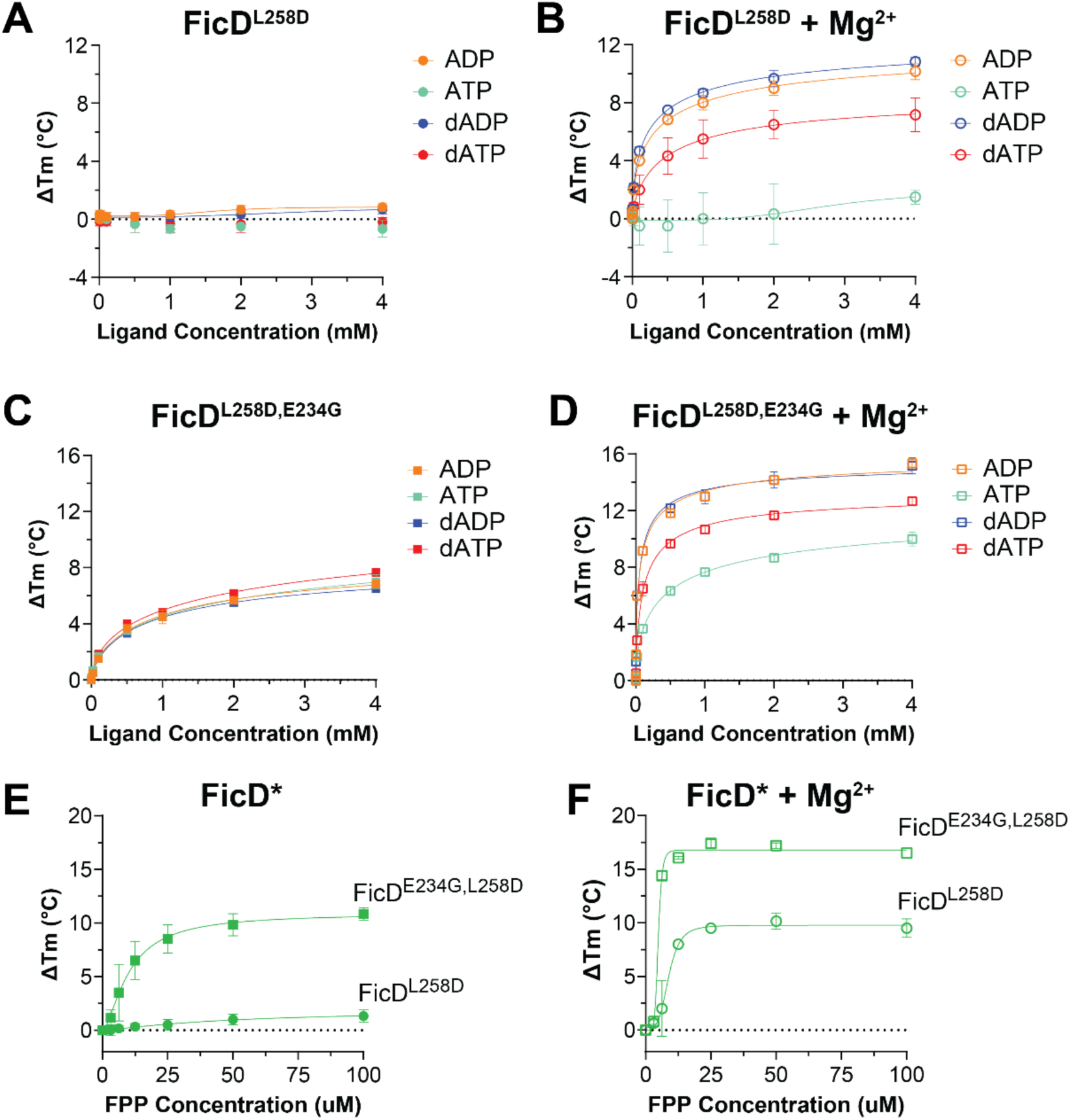
Mg^2+^ increases nucleotide and FPP binding to FicD^L258D^ and FicD^E234G, L258D^. **(A-D)** Dose response FicD melting by DSF with nucleotides: ADP (orange), ATP (teal), dADP (blue), and dATP (red). **(A)** FicD^L258D^ in the absence or **(B)** presence of 5mM MgCl_2_ and **(C)** FicD^E234G, L258D^ in the absence **(D)** or presence of 5mM MgCl_2_. **(E and F)** FPP-induced DSF melting of Fic^L258D^ and FicD^E234G, L258D^ **(E)** in the absence or **(F)** presence of 5mM MgCl_2_. Error bars for all curves represent three independent experiments.

Interestingly in an alternate FicD variant, FicD^E234G, L258D^, all nucleotides were able to bind in the absence of MgCl_2_ (**Fig 4C**). The addition of 5mM MgCl_2_ further enhanced binding of these nucleotides (**Fig 4D**). To assess if the pyrophosphate of FPP could coordinate Mg^2+^ in a manner analogous to that of the nucleotides, we asked if FPP binding also required Mg^2+^. Interestingly, the pattern of FPP binding to the FicD variants was like that of ADP, dADP, and dATP. In the absence of MgCl_2_, FPP did not bind to FicD^L258D^; whereas it bound to FicD^E234G, L258D^ (**Fig 4E**). In the presence of 5mM MgCl_2_, FPP bound to both FicD^L258D^ and FicD^E234G,L258D^ (**Fig 4F**). These similarities in the Mg^2+^ requirement for FPP and nucleotide binding suggested that FPP could adopt a nucleotide-like binding mode in the active site of FicD.

### Structural analysis reveals FPP binding to the active site of FicD

To reveal the mechanism of FicD inhibition by FPP, we leveraged the FicD^E234G^ mutant for structural studies. We soaked FPP into crystals of the FicD^E234G^ variant and solved the ligand-bound structure. The 2.6 Å resolution ligand bound structure revealed binding of FPP to FicD active site. The diphosphate group of FPP closely overlapped with the ADP diphosphate when superimposed with a FicD^E234G^-ADP structure (**Fig 5A**, left panel), and the hydrophobic isoprenyl groups of FPP overlap with the nucleoside in the active site binding pocket (**Fig 5A**, right panel). The FPP diphosphate binds directly to the side chains of N369, R371, with the sidechain of D367 coordinating the Mg^2+^ (**Fig. 5B**). These interactions mimic those observed between ADP/ATP and FicD (26). Total mass analysis of FicD^WT^ or FicD^E234G^ incubated with FPP demonstrated no mass shift associated with a putative FPP auto-modification (**Fig S6A and S6B**), consistent with non-catalytic positioning of the FPP diphosphates. As previously noted, only GPP and FPP could bind to and inhibit FicD, but analogs of the isoprenoid-pyrophosphate metabolites (geraniol, GMP, farnesol, FMP, DMAPP, and MVAPP) could not (**Fig 2F**). Together, these observations indicate both the pyrophosphate head and the long isoprenoid tail of GPP/FPP contribute to the binding with FicD. Importantly, this also implies potential for the FicD pocket to be targeted when designing diverse inhibitor molecules that use diphosphate as a polar head.

**Figure 5.**
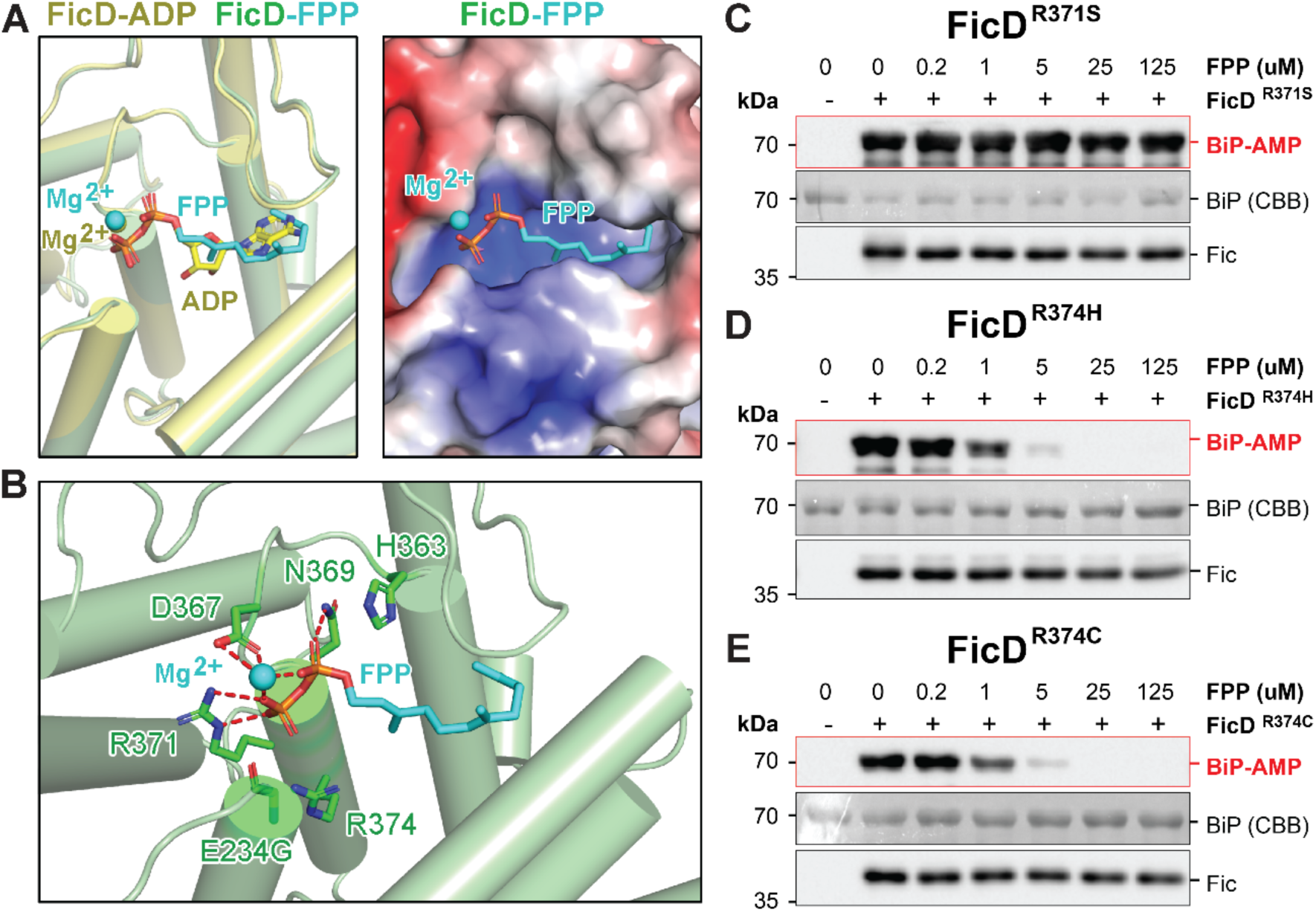
Crystal structure of FPP inhibitor bound to FicD active site. **(A)** FicD-FPP structure in green aligned with FicD-ADP in yellow (PDB: 4U0S) shows FPP (cyan stick) and Mg^2+^ (cyan sphere) binding the active site like Mg^2+^ADP (yellow sphere and stick). **(B)** FPP (cyan stick) and Mg^2+^ (cyan sphere) in the pocket of FicD shown as APBS electrostatic surface with positive areas in blue, negative in red, and neutral in white (contour level: +/- 5 *k*BT/e). **(C)** Mg^2+^FPP (cyan) forms hydrogen bonds (dashed red lines) with the side chains of D367, N369, and R371 near the catalytic H363, but does not interact with R374. **(D-F)** AMPylation assays of Δ26hBiP^T229A^ in the presence of the indicated concentration of FPP using the pathogenic human variants **(D)** FicD^R371S^, **(E)** FicD^R374H^ and **(F)** FicD^R374C^. AMPylated BiP and Fic were quantified by immunodetection, and total BiP was quantified by CBB.

### Interaction of Farnesyl-pyrophosphate with pathogenic variants of FicD

Three FicD variants have been implicated in human disease. All three variants, FicD^R371S^, FicD^R374H^, and FicD^R374C^, occur at conserved active site FicD residues. Their mutation confers total loss of deAMPylation activity while also lowering AMPylation activity (39–42). The sidechain of FicD R371 directly facilitates binding of the β-phosphates of FPP and ADP (**Fig. 5B**). Given this interaction, we predicted R371 to be critical for facilitating FPP binding and its inhibition of FicD activity. Indeed, the FicD^R371S^ variant exhibited a total loss of FPP inhibition of AMPylation activity (**Fig 5C**). Additionally, the addition of 5mM MgCl_2_ binding did not confer FPP binding to the FicD^R371S^ variant at any concentration tested (**Fig S7A and S7B**). These results confirm that loss of FPP-mediated AMPylation inhibition was due to loss of FPP binding conferred by the FicD^R371S^ mutation.

Unlike the R371 sidechain, FicD R374 does not directly interact with FPP (**Fig 5B**). Instead, R374 either coordinates the gamma phosphate of ATP in the AMPylation-competent state, or it forms the autoinhibitory salt bridge with E234 in the deAMPylation competent state of the enzyme (49). Therefore, mutation of R374 to His or Cys should break the autoinhibitory salt bridge, mirroring the E234G mutation that confers constitutive AMPylation. The R374H and R374C variants were not predicted to disrupt FPP binding because the gamma phosphate that R374 interacts with is missing in FPP. Indeed, using DSF, we observe shifts in the melting curves of the R374H and R374C variants upon addition of FPP. Interestingly, FPP caused shifts in the melting curves of the R374H and R374C mutants in the absence of Mg^2+^ (**Fig S7C and S7D**), though the addition of Mg^2+^ enhanced the binding (**Fig S7E and S7F**). These observations are consistent with R374 being required for the autoinhibitory salt bridge. In activity assays, we observe FPP inhibition of FicD^R374H^ and FicD^R374C^ mediated AMPylation (**Fig 5D and 5E**). Together, these data support our structural findings that FPP engages conserved active site residues analogous to ADP and highlights active site differences in the pathogenic variants that can be targeted for different inhibition profiles.

## Discussion

Metazoan FicD utilizes a bifunctional Fido domain to catalyze AMPylation and deAMPylation of the essential HSP70 chaperone BiP (23, 24). The bifunctional activity of FicD promotes adaptive engagement of UPR stress signaling by fine tuning ER chaperone capacity (21). While it is known that FicD’s bifunctional activity is influenced by ER stress, the in-vivo signaling inputs that regulate FicD activity remain unclear. Therefore, we utilize unbiased high-throughput screening (MIDAS) to identify small molecule metabolites that might regulate FicD. We identify isoprenoid diphosphates GPP and FPP as potent and specific inhibitors of FicD-mediated AMPylation and deAMPylation. Using structural biology, we observed FPP bound to the active site of FicD analogous to binding of ADP, which engages a conserved active site residue R371.

To investigate the possibility of additional physiological FicD ligands, MIDAS was selected for its small molecule library composed of naturally occurring metabolites. Our MIDAS screen reinforces the known FicD’s specificity for FicD binding adenine nucleotides. The thyroid hormone metabolites 3,3’-diiodothyronine and 3’-monoiodo-L-thyronine were also significantly enriched in the MIDAS screen (**Fig 1C**). This observation mirrors the results of a previous drug screen with FicD that identified liothyronine as a FicD inhibitor (45). However, it is notable that no other classes of small molecules were identified as hits across all of the tested FicD constructs. This observation implies that the natural ligands that interact with FicD occupy a limited chemical space.

Using MIDAS, we identified GPP/FPP as new PMIs for FicD, and these metabolites inhibited both AMPylation or deAMPylation activities. It is possible that independent bifunctional activities of FicD are not regulated by a small molecule metabolite but is instead regulated by another unknown cellular factor or modification, similar to other AMPylators (50). For example, FicD activity may be indirectly dictated by alterations to BiP chaperone cycling. FicD specifically recognizes the ATP-bound form of BiP (22, 51). BiP’s nucleotide binding is dynamic and regulated by several factors, such as ER Ca^2+^ levels, rate of ER ATP/ADP exchange, and action of nucleotide exchange factors, which are all affected by ER stress. As such, the possibility that FicD activity is correlated to substrate abundance has been proposed (31). As FicD is an ER membrane protein, its oligomerization activity switch may also be influenced by the membrane bilayer.

Given the localization of metazoan FicD active site to the ER lumen, the biological relevance of FPP/GPP binding and inhibition of BiP AMPylation and deAMPylation is not apparent. No known GPP/FPP utilizing enzymes localize to the luminal side of the ER membrane. Instead, the GPP/FPP inhibitors are synthesized on the cytosolic face of the ER, where they are not compatible with binding to the FicD active site. However, disregulation of lipid synthesis and UPR are know to play roles in type 2 diabetes, liver disease and neurodegenerative disorders (52). Therefore it is tempting to speculate that unknown roles for GPP/FPP exist in th ER lumen and, therefore, could affect Fic activity. The UPR and mevalonate pathways are intrinsically linked through cholesterol and lipid synthesis, thus regulation of FicD activity by GPP/FPP could be an evolved regulatory mechanism.

Another regulatory possibility is with isoprenoids and glycosylation. Isoprenoids serve as precursors for a longer dolichol phosphate lipid carrier essential for N-linked protein glycosylation in the ER lumen (48). After N-linked glycosylation of ER proteins, the dolichol is released in its diphosphate form (Dol-PP) on the lumenal side of the ER membrane (52). While it is tempting to speculate that FicD activity should respond to N-linked glycosylation of newly synthesized ER proteins, whether the hydrophobic Dol-PP could partition between the membrane and the FicD active site remains to be determined. Biochemically testing this questions is challenging due to the solubilty of Dol-PP.

The K_d_ of FPP binding to FicD was 60nM. This relatively tight binding is over an order of magnitude stronger than the K_d_ for ADP (26). Despite this notable inhibition of the human FicD enzyme, the activities of two constitutively active bacterial Fido proteins, AvrB and VopS, were not inhibited by FPP (**FIG S5**). However, the Fido domains of AvrB and VopS are structurally and functionally distinct from FicD (20, 25, 30). Neither are intramolecularly autoinhibited nor are they bifunctional. Indeed, it is possible that GPP/FPP binding requires autoinhibition and that other autoinhibited Fido proteins interact with GPP/FPP. Approximately 90% of all AMPylating Fic domains are predicted to be intrinsically inhibited and bifunctional, a majority of which are in bacteria (32). Like metazoan FicD, these enzymes play a housekeeping role, modifying DNA gyrases and topoisomerases to tune bacterial stress signaling (53, 54). The bacterial Fido domain proteins are localized to the cytoplasm where terpenoid biosynthesis from FPP intermediates is performed (55, 56). Therefore, our findings may simply reflect an evolutionary link that remains between bacterial housekeeping Fido domains that function in stress signaling and their metazoan counterparts that retain intramolecular autoinhibition.

FicD^R371S^, FicD^R374H^, and FicD^R374C^ variants are implicated in human disease. FicD^R371S^ causes a rare form of neonatal diabetes with severe neurodevelopmental impairment (39). FicD^R374H^ and FicD^R374C^ are implicated in a progressive motor neuron disease and increased risk of diabetes in adulthood (41, 42). In all three variants, loss of BiP deAMPylation is central to the mechanism of disease (39, 41, 42) Active site changes in the pathogenic variants contribute to FPP binding. FicD^R374H^ and FicD^R374C^ retain interaction with FPP and their AMPylation activity is inhibited by the metabolite (**Fig S7B-C, Fig 5D-E**). Conversely, AMPylation activity of the FicD^R371S^ variant lacking FPP binding was not inhibited (**Fig 5A**). These results are consistant with the contribution of R371 and R374 to FPP binding observed in our crystal structure (**Fig 5B**).

A preclinical FicD^R371S^ mouse model demonstrated progressive loss of pancreatic islet organization with defects in insulin secretion (40), which mirrors phenotypes in FicD^R371S^ homozygote children (39). In the FicD^R374H^ and FicD^R374C^ variants, loss of intramolecular autoinhibition confers loss of deAMPylation activity. This mechanism mirrors the biochemically well-characterized FicD^E234G^, which is homozygous lethal in both flies (24) and mice (38, 43). The E234G mutation confers enhanced Mg^2+^ independent binding of FPP, which was also observed in the FicD^R374H^ and FicD^R374C^ mutants. Therefore, our crystal structure of FPP bound to FicD may provide an experimentally grounded foundation for future medicinal chemistry efforts to develop inhibitors to treat pathologic FicD AMPylation caused by the FicD^R374H^ and FicD^R374C^ mutations.

In summary, we characterize the interaction of FPP with FicD in precise biochemical and structural detail. The FPP specificity and its potency of action in inhibiting FicD is particularly striking. FPP binds to the FicD active site with over an order of magnitude higher affinity than canonical ligands. The observed interactome of FicD with natural ligands was also markedly small due to our triage strategy. These included nucleotides, thyroid hormone intermediates and GPP/FPP. In addition, the interaction with GPP/FPP may be exclusive to intramolecularly autoinhibited Fic proteins. Together, these observations implicate a physiological role for isoprenoid metabolites and FicD activity. Finally, we define dADP and dATP as novel FicD ligands, potentially expanding the repertoire of cosubstrates utilized by the Fido domain. Together, our work provides novel insight into ligands that can bind to the active site of the FicD, while revealing catalytic specificity for human disease variants.

## Materials and Methods

Detailed descriptions of the experimental methods are provided in SI Appendix. These include reagents, high-throughput screening, data analysis of screening data, cloning of genes, protein purification, in-vitro activity and thermal shift assays, protein crystallography, and isothermal titration calorimetry.

## Acknowledgments

We thank Chad Braudingham and Scott Tso at the UT Southwestern Macromolecular Biophysics core for their assistance with ITC experiments. We thank David Hein for assistance in the thermal shift data analysis. We thank Andy Lemoff at the UT Southwestern Proteomics Core for assistance with LC/MS analysis of intact proteins. We thank the Structural Biology Lab at UT Southwestern Medical Center for support with X-ray crystallographic studies. We thank Aymelt Itzen at the University Medical Center Hamburg Eppendorf, Germany for providing monoclonal AMPylation antibodies. We thank the Orth lab members for discussions and editing.

Results shown in this report are derived from work performed at the Berkeley Center for Structural Biology at the Advanced Light Source (ALS). The Berkeley Center for Structural Biology is supported by the Howard Hughes Medical Institute, Participating Research Team members, and the National Institutes of Health, National Institute of General Medical Sciences, ALS-ENABLE grant P30 GM124169. The ALS is a Department of Energy Office of Science User Facility under Contract No. DE-AC02-05CH11231. The Pilatus detector on beamline 2.0.1 was funded under NIH grant S10OD021832. The Pilatus detector on beamline 5.0.1 was funded under NIH grant S10OD026941. KO is a WW Caruth, Jr. Biomedical Scholar with an Earl A Forsythe Chair in Biomedical Science. Funding Sources: Welch Foundation grant I-1561 (KO); Once Upon a Time…Foundation (KO); the National Institutes of Health Grant R35 GM134945 (KO) and R35 GM131854 (JR); the National Institutes of Health T32 Chemistry-Biology Interface Training Grant T32GM127216 (AMB); Chan Zuckerberg Initiative, project number: 10070558 / 51006865 (KGH). The funders had no role in study design, data collection and interpretation, or the decision to submit the work for publication.

## Data and materials availability

The atomic coordinates of the corresponding crystal structure are deposited in the Protein Data Bank with accession code 9YZ5 (extended ID pdb_00009YZ5).

## References

1. L. M. Hendershot, T. M. Buck, J. L. Brodsky, The Essential Functions of Molecular Chaperones and Folding Enzymes in Maintaining Endoplasmic Reticulum Homeostasis. J Mol Biol 436, 168418 (2024).

2. I. Braakman, D. N. Hebert, Protein folding in the endoplasmic reticulum. Cold Spring Harb Perspect Biol 5, a013201 (2013).

3. V. Gonzalez-Teuber et al., Small Molecules to Improve ER Proteostasis in Disease. Trends Pharmacol Sci 40, 684–695 (2019).

4. X. Chen, C. Shi, M. He, S. Xiong, X. Xia, Endoplasmic reticulum stress: molecular mechanism and therapeutic targets. Signal Transduct Target Ther 8, 352 (2023).

5. C. Hetz, K. Zhang, R. J. Kaufman, Mechanisms, regulation and functions of the unfolded protein response. Nat Rev Mol Cell Biol 21, 421–438 (2020).

6. C. Lebeaupin, J. Yong, R. J. Kaufman, The Impact of the ER Unfolded Protein Response on Cancer Initiation and Progression: Therapeutic Implications. Adv Exp Med Biol 1243, 113–131 (2020).

7. B. M. Gardner, D. Pincus, K. Gotthardt, C. M. Gallagher, P. Walter, Endoplasmic reticulum stress sensing in the unfolded protein response. Cold Spring Harb Perspect Biol 5, a013169 (2013).

8. D. Pincus et al., BiP binding to the ER-stress sensor Ire1 tunes the homeostatic behavior of the unfolded protein response. PLoS Biol 8, e1000415 (2010).

9. A. Bertolotti, Y. Zhang, L. M. Hendershot, H. P. Harding, D. Ron, Dynamic interaction of BiP and ER stress transducers in the unfolded-protein response. Nat Cell Biol 2, 326–332 (2000).

10. J. Shen, X. Chen, L. Hendershot, R. Prywes, ER stress regulation of ATF6 localization by dissociation of BiP/GRP78 binding and unmasking of Golgi localization signals. Dev Cell 3, 99–111 (2002).

11. A. Bakunts et al., Ratiometric sensing of BiP-client versus BiP levels by the unfolded protein response determines its signaling amplitude. Elife 6 (2017).

12. M. Vitale et al., Inadequate BiP availability defines endoplasmic reticulum stress. Elife 8 (2019).

13. D. Acosta-Alvear, J. M. Harnoss, P. Walter, A. Ashkenazi, Homeostasis control in health and disease by the unfolded protein response. Nat Rev Mol Cell Biol 26, 193–212 (2025).

14. L. Plate, R. L. Wiseman, Regulating Secretory Proteostasis through the Unfolded Protein Response: From Function to Therapy. Trends Cell Biol 27, 722–737 (2017).

15. S. J. Marciniak, J. E. Chambers, D. Ron, Pharmacological targeting of endoplasmic reticulum stress in disease. Nat Rev Drug Discov 21, 115–140 (2022).

16. C. Hetz, J. M. Axten, J. B. Patterson, Pharmacological targeting of the unfolded protein response for disease intervention. Nat Chem Biol 15, 764–775 (2019).

17. H. Ham et al., Unfolded protein response-regulated Drosophila Fic (dFic) protein reversibly AMPylates BiP chaperone during endoplasmic reticulum homeostasis. J Biol Chem 289, 36059–36069 (2014).

18. A. Sanyal et al., A novel link between Fic (filamentation induced by cAMP)-mediated adenylylation/AMPylation and the unfolded protein response. J Biol Chem 290, 8482–8499 (2015).

19. S. Preissler et al., AMPylation matches BiP activity to client protein load in the endoplasmic reticulum. Elife 4, e12621 (2015).

20. L. N. Kinch, M. L. Yarbrough, K. Orth, N. V. Grishin, Fido, a novel AMPylation domain common to fic, doc, and AvrB. PLoS One 4, e5818 (2009).

21. A. K. Casey et al., Fic-mediated AMPylation tempers the unfolded protein response during physiological stress. Proc Natl Acad Sci U S A 119, e2208317119 (2022).

22. S. Preissler et al., AMPylation targets the rate-limiting step of BiP’s ATPase cycle for its functional inactivation. Elife 6 (2017).

23. S. Preissler, C. Rato, L. Perera, V. Saudek, D. Ron, FICD acts bifunctionally to AMPylate and de-AMPylate the endoplasmic reticulum chaperone BiP. Nat Struct Mol Biol 24, 23–29 (2017).

24. A. K. Casey et al., Fic-mediated deAMPylation is not dependent on homodimerization and rescues toxic AMPylation in flies. J Biol Chem 292, 21193–21204 (2017).

25. M. L. Yarbrough et al., AMPylation of Rho GTPases by Vibrio VopS disrupts effector binding and downstream signaling. Science 323, 269–272 (2009).

26. T. D. Bunney et al., Crystal structure of the human, FIC-domain containing protein HYPE and implications for its functions. Structure 22, 1831–1843 (2014).

27. B. Gulen, A. Itzen, Revisiting AMPylation through the lens of Fic enzymes. Trends Microbiol 30, 350–363 (2022).

28. P. Engel et al., Adenylylation control by intra- or intermolecular active-site obstruction in Fic proteins. Nature 482, 107–110 (2012).

29. A. Goepfert, F. V. Stanger, C. Dehio, T. Schirmer, Conserved inhibitory mechanism and competent ATP binding mode for adenylyltransferases with Fic fold. PLoS One 8, e64901 (2013).

30. W. Peng et al., Pseudomonas effector AvrB is a glycosyltransferase that rhamnosylates plant guardee protein RIN4. Sci Adv 10, eadd5108 (2024).

31. L. A. Perera et al., An oligomeric state-dependent switch in the ER enzyme FICD regulates AMPylation and deAMPylation of BiP. Embo j 38, e102177 (2019).

32. F. V. Stanger et al., Intrinsic regulation of FIC-domain AMP-transferases by oligomerization and automodification. Proc Natl Acad Sci U S A 113, E529–537 (2016).

33. S. Veyron et al., A Ca(2+)-regulated deAMPylation switch in human and bacterial FIC proteins. Nat Commun 10, 1142 (2019).

34. M. Rahman et al., Visual neurotransmission in Drosophila requires expression of Fic in glial capitate projections. Nat Neurosci 15, 871–875 (2012).

35. A. G. Lobato et al., Loss of Fic causes progressive neurodegeneration in a Drosophila model of hereditary spastic paraplegia. Biochim Biophys Acta Mol Basis Dis 1870, 167348 (2024).

36. A. K. Casey et al., FicD regulates adaptation to the unfolded protein response in the murine liver. Biochimie 225, 114–124 (2024).

37. S. M. Lacy et al., FICD (FIC Domain Protein Adenylyl Transferase) Deficiency Protects Mice From Hypertrophy-Induced Heart Failure and Promotes BiP (Binding Immunoglobulin Protein) -Mediated Activation of the Unfolded Protein Response and Endoplasmic Reticulum-Selective Autophagy in Cardiomyocytes. J Am Heart Assoc 14, e040192 (2025).

38. K. M. Van Pelt, M. C. Truttmann, Loss of FIC-1-mediated AMPylation activates the UPRER and upregulates cytosolic HSP70 chaperones to suppress polyglutamine toxicity. PLoS Genet 21, e1011723 (2025).

39. L. A. Perera et al., Infancy-onset diabetes caused by de-regulated AMPylation of the human endoplasmic reticulum chaperone BiP. EMBO Mol Med 15, e16491 (2023).

40. A. K. Casey et al., Pre-clinical model of dysregulated FicD AMPylation causes diabetes by disrupting pancreatic endocrine homeostasis. Mol Metab 95, 102120 (2025).

41. A. P. Rebelo et al., BiP inactivation due to loss of the deAMPylation function of FICD causes a motor neuron disease. Genet Med 24, 2487–2500 (2022).

42. A. P. Rebelo et al., FIC Domain Protein Adenylyltransferase (FICD)-Related Complex Hereditary Spastic Paraplegia with Diabetes Mellitus. Mov Disord Clin Pract 12, 1193–1195 (2025).

43. N. McCaul et al., Deletion of mFICD AMPylase alters cytokine secretion and affects visual short-term learning in vivo. J Biol Chem 297, 100991 (2021).

44. K. G. Hicks et al., Protein-metabolite interactomics of carbohydrate metabolism reveal regulation of lactate dehydrogenase. Science 379, 996–1003 (2023).

45. B. K. Chatterjee et al., Small-Molecule FICD Inhibitors Suppress Endogenous and Pathologic FICD-Mediated Protein AMPylation. ACS Chem Biol 20, 880–895 (2025).

46. A. B. Uceda, L. Mariño, R. Casasnovas, M. Adrover, An overview on glycation: molecular mechanisms, impact on proteins, pathogenesis, and inhibition. Biophys Rev 16, 189–218 (2024).

47. M. Kanehisa, Y. Sato, M. Kawashima, M. Furumichi, M. Tanabe, KEGG as a reference resource for gene and protein annotation. Nucleic Acids Research 44, D457–D462 (2015).

48. M. B. Jones, J. N. Rosenberg, M. J. Betenbaugh, S. S. Krag, Structure and synthesis of polyisoprenoids used in N-glycosylation across the three domains of life. Biochim Biophys Acta 1790, 485–494 (2009).

49. L. A. Perera et al., Structures of a deAMPylation complex rationalise the switch between antagonistic catalytic activities of FICD. Nat Commun 12, 5004 (2021).

50. A. K. Casey, K. Orth, Enzymes Involved in AMPylation and deAMPylation. Chem Rev 118, 1199–1215 (2018).

51. J. Fauser et al., Specificity of AMPylation of the human chaperone BiP is mediated by TPR motifs of FICD. Nat Commun 12, 2426 (2021).

52. W. Bialek et al., The lipid side of unfolded protein response. Biochim Biophys Acta Mol Cell Biol Lipids 1869, 159515 (2024).

53. A. Harms et al., Adenylylation of Gyrase and Topo IV by FicT Toxins Disrupts Bacterial DNA Topology. Cell Rep 12, 1497–1507 (2015).

54. C. Lu, E. S. Nakayasu, L. Q. Zhang, Z. Q. Luo, Identification of Fic-1 as an enzyme that inhibits bacterial DNA replication by AMPylating GyrB, promoting filament formation. Sci Signal 9, ra11 (2016).

55. E. J. N. Helfrich, G. M. Lin, C. A. Voigt, J. Clardy, Bacterial terpene biosynthesis: challenges and opportunities for pathway engineering. Beilstein J Org Chem 15, 2889–2906 (2019).

56. J. Pérez-Gil, M. Rodríguez-Concepción, Metabolic plasticity for isoprenoid biosynthesis in bacteria. Biochem J 452, 19–25 (2013).

