## Supplementary material for "Inhibition of FicD-mediated AMPylation and deAMPylation by Isoprenoid Diphosphates": supp file

**This PDF file includes:**

Supporting text
Figures S1 to S7
Tables S1 to S6
Legends for Datasets S1 to S4
SI References

**Other supporting materials for this manuscript include the following:**

Datasets S1 to S4

### Supporting Information Text

### SI Materials and Methods

*Chemicals and antibodies*

dADP, dATP, Geraniol, Geranyl-pyrophosphate, Farnesol, Farnesyl-pyrophosphate, Dimethylallyl-pyrophosphate, and Mevalonate-pyrophosphate were purchased from MedChemExpress (cat nos. HY-W010854, HY-W013098, HY-N6952R, HY-114295A, HY-Y0248AR, HY-113037B, HY-130573A, and HY-N9474). ADP, ATP, Isopententyl-pyrophsophate, Geranyl-monophosphate and Farnesyl-monophosphate were purchased from Sigma (Cat nos. A7699, A2754, I0503, 56901 and 91356). Geraniol and Farnesol were dissolved in 100% DMSO. dADP, dATP, ADP, ATP, Geranyl-pyrophosphate, Farnesyl-pyrophosphate, Dimethylallyl-pyrophosphate, and Mevalonate-pyrophosphate were dissolved in H2O. Farnesyl-monophosphate was dissolved in 100% methanol. Geranyl-monophosphate was dissolved in Methanol:DMSO:H2O 1:1:4.

Antibodies used in this study are as follows: α-AMP (17G6) (1) (gift from Aymelt Itzen), α-Fic (custom antibody from Thermo/Fisher against GDVRPFIRFIAKCTET peptide), α-Rin4-T166-Rha (2), α-GST (ProteinTech cat no. 10000-0-AP), goat-α-rabbit-HRP (Amersham cat no. NA934), goat α-mouse-HRP (Abcam cat no. ab205719 or Sigma cat no. A9917).

*Cloning of constructs*

FicD and BiP genes were amplified from human cDNA clones purchased from the UT Southwestern McDermott Center for Human Growth and Development with Thermo Scientific with Phusion High-Fidelity DNA Polymerase (cat no. F530) (**Table S1**). Amplicons were digested with Thermo Scientific FastDigest BamHI (cat no. FD0054) and NotI (cat no. FD0595) prior to cleanup with BioBasic PCR Products Purification kit (cat no. BS365). pET28a-SUMO and pET28A vectors were digested with the same restriction enzymes prior to gel purification with BioBasic Gel Extraction Kit (cat no. BS654). The FicD insert was ligated into the ppSumo vector and the BiP insert was ligated into the pET28A vector with Thermo Scientific T4 DNA Ligase (cat no. EL0011). Ligation products were then transformed into Mach 1 E. coli (Invitrogen cat no. C862003).

Site directed mutagenesis of FicD and BiP and truncation of FicD c-terminal residues was performed by inverse PCR using primers in **Table S1** with either Thermo Scientific with Phusion High-Fidelity DNA Polymerase (cat no. F530) or NEB Q5 DNA Polymerase (cat no. M0491). Amplicons were treated with Thermo Scientific FastDigest DpnI (FD1704) prior to cleanup. Plasmid circularization was then performed with either NEB T4 Polynucleotide kinase (cat no. M0201) and Thermo Scientific T4 DNA Ligase (cat no. EL0011) or NEB KLD enzyme mix (cat no. M0554S) prior to transformation into Mach 1 *E. coli* (Invitrogen cat no. C862003).

Transformants were screened with Thermo Scientific DreamTaq DNA Polymerase (cat no. K1081) and insert-positive colonies were miniprepped with BioBasic Plasmid DNA Miniprep Kit (cat no. BS614). Plasmids were sequenced with Genewiz Sanger sequencing or PlasmidSaurus whole-plasmid sequencing.

*Protein expression and purification*

Wild-type FicD (residues 105-458) and all mutants were expressed as N-terminal 6xHis-Sumo fusions. One construct of FicD$^{E234G}$ – encompassing residues 105-433 - was used exclusively for crystallography. BiP$^{T229A}$ (residues 27-654) was expressed as an 6xHis-Thrombin-T7 n-terminal fusion. Expression plasmids were transformed into Rosetta(DE3) *E. coli*. A single colony transformant was then inoculated into a 5mL of 2X YT with 30ug/mL Kanamycin and 10ug/mL Chloramphenicol. The starter culture was nutated at 37°C for ~4 hours and then transferred into 250-500mL of 2X YT with 30µg/mL Kanamycin and 10µg/mL Chloramphenicol. The culture was nutated at 37°C until OD600 reached 0.6-0.8 and then chilled to induction temperature. Expression of all FicD constructs was induced with 50µM IPTG overnight (approximately 18 hours) at 16C-18°C. Expression of BiP$^{T229A}$ was induced with 400µM IPTG overnight (approximately 18 hours) at

22°C. Induced cultures were then harvested by centrifugation at 8000G for 15 minutes at 4°C and stored at -80°C.

Bacterial pellets were resuspended in 30-50mL of 50mM Tris pH 7.4/150mM NaCl/10% glycerol with one Roche cOmplete EDTA-Free Protease inhibitor cocktail tablet (cat no. COEDTAF-RO). Triton-X100 to a final concentration of 0.2%, Chicken-egg lysozyme to a final concentration of 0.4ug/mL, and 2-5uL of Millipore Benzonase Nuclease (cat no. 70664) were added to bacterial suspension prior to sonication at 40-50% amplitude for 1-3 minutes in pulses of 5 seconds on and 30 seconds off on ice with a Sonics VCX 750 Ultrasonicator. Bacterial lysate was then clarified by centrifugation at 30,000 x g for 30 minutes at 4°C.

Clarified lysate was rotated with Qiagen NiNTA agarose beads (cat no. 30210) for 1-2 hours at 4°C. The beads were then transferred to a 15mL conical vial, resuspended in 10-15mL wash buffer (50mM Tris pH 7.4/150mM NaCl/10% glycerol/0.2% Triton-X 100/10mM imidazole), and spun down at 1000G for 1 minute at 4°C, after which the supernatant was aspirated from the beads. This washing step was repeated 2-4 times to rinse bacterial lysate from the beads prior to on-column washing. The beads were then transferred to BioRad Econopac 20mL bed volume column (cat no. 7321010) and washed with 10-15mL of wash buffer (repeated at least three times), 10mL of ATP wash (50mM Tris pH 7.4/300mM KCl/10mM MgCl2/500uM ATP, repeated once), and finally with 10-15mL wash buffer without Triton-X 100 (repeated at least three times). Protein was eluted in 500µL-1mL elution buffer (50mM Tris pH 7.5/150mM NaCl/10% glycerol/0.5% CHAPS). For all Fic constructs, elutions were supplemented with DTT and EDTA to a final concentration of 1mM and ULP-protease (gift from Vincent Tagliabracci) and rotated at 4°C to remove the 6xHis-Sumo tag.

Elutions were clarified either by centrifugation at 21,000G for 15 minutes at 4C or with MilliPore UltraFree 0.22uM centrifugal filter (cat no. UFC30GV0S) prior to gel filtration with 50mM Tris pH 7.4/150mM NaCl/10% glycerol in a Cytiva HiLoad 16/600 Superdex 200 pg column at a flow rate of 1mL/minute at 4°C. Impurity-free elution fractions were then combined and concentrated in Millipore Amicon centrifugal concentrators. All FicD constructs were concentrated in 30 kDa MWCO concentrators (cat no. UFC9030) and BiP was concentrated in 50kDa MWCO concentrator (cat no. UFC9050). Concentrated proteins were then clarified either by centrifugation at 21,000G for 15 minutes at 4°C or with a MilliPore UltraFree 0.22uM centrifugal filter. Finally, DTT was added to clarified protein to a final concentration of 1mM. Proteins were aliquoted, flash frozen in liquid nitrogen, and then stored at -80C.

*MIDAS Metabolite Library*
The MIDAS metabolite library was constructed similarly to what was previously described (3). Briefly, all metabolites used in this study were purchased from Sigma-Aldrich, Cayman Chemicals, Avanti Polar Lipids, Enamine, Combi-Blocks, Inc, or custom sourced from Molport. Metabolites were solvated to 10 mM in molecular grade water (Millipore Sigma W4502) or DMSO (Millipore Sigma D1435) and, where necessary to increase solubility, titrated with acid or base. The MIDAS metabolite library was arrayed 1 mL per well in 96-deep well storage plates (Greiner 780280), and sealed with aluminum foil seals (Greiner 676090), and stored at -80°C. When working stocks were needed, metabolites were moved from deep well storage plates and arrayed, 50 µL per well, across multiple 384-well small volume storage plates (Greiner 781280), sealed with aluminum foil seals (Greiner 676090), and stored at -80°C. Metabolite library management, manipulation, and pooling was conducted with a Beckman Coulter Biomek NXp SPAN-8 liquid handling robot.

*MIDAS Equilibrium Dialysis*
The day of MIDAS analysis, the 384-small volume working stock plates of the MIDAS metabolite library were defrosted at 37°C for 10 minutes. All metabolites were combined into one pool at a final concentration of 5 µM in 25 mM Hepes pH 7.4 (Fisher Scientific BP310), and 100 mM NaCl (Fisher Scientific BP358) and pH adjusted with sodium hydroxide (Fisher Scientific 424330025). 50 µL of each of 360 µM $\Delta104hFic^{E234G}$; 603 µM $\Delta104hFic^{L258D}$; 447 µM $\Delta104hFic^{H363A}$; 496 µM $\Delta104hFic^{WT}$; 456 µM $\Delta104hFic^{L258D}$; and 224 µM $\Delta104hFic^{H363A}$ protein in buffer containing 25 mM Hepes pH 7.4 (Fisher Scientific BP310), and 100 mM NaCl (Fisher Scientific BP358) was arrayed,

in triplicate, in the red chambers of a 8 kDa Rapid Equilibrium Dialysis Device Insert (Thermo Fisher Scientific 89809) in a Rapid Equilibrium Dialysis Device Reusable Base Plate (Thermo Fisher Scientific 89811). 300 μL of the metabolite pool was aliquoted into the white chamber of each insert and the plate was sealed with an aluminum foil seal (Greiner 676090). The loaded Rapid Equilibrium Dialysis Device plate was placed at 4°C on a microplate shaker at 400 rpm and incubated for 24 hours. Post dialysis, protein and metabolite chamber dialysates were retrieved, snap frozen in liquid nitrogen and placed at -80°C. When ready for metabolomic analyses, protein and metabolite chamber samples were thawed and diluted 1:4 in ice-cold 100% methanol (Thermo Fisher Scientific 047192) to denature protein, incubated at 30 minutes at -20°C, and centrifuged at 21,100 x g for 30 minutes to remove precipitated protein. Processed protein and metabolite chamber samples were retrieved and arrayed across a 384-well microplate (Greiner 781281) and heat sealed with heat sealing aluminum foil (Thermo Fisher Scientific AB-0757) using an ALPS 50 V-Manual Heat Sealer (Thermo Fisher Scientific AB-1443A) at 160°C for 2.5 seconds. The sealed sample plates were placed at 4°C for analysis by liquid chromatography-mass spectrometry (LC-MS) metabolomics.

*MIDAS LC-MS metabolomics*
The MIDAS LC-MS platform was composed of an electrospray ionization (ESI), quadrupole time-of-flight (QTOF) LC-MS system equipped for hydrophilic interaction liquid chromatography (HILIC) metabolomics. The LC-MS system was composed of a Shimadzu Nexera HPLC system equipped with binary LC-40DXR pumps (A and B), DGU-405 degassing unit, CBM-40 system controller, CTO-40C column oven (maintained at 40°C), and a SIL-40C XR autosampler (maintained at 4°C) coupled to a SCIEX X500R ESI-QTOF MS operated in positive and negative mode. Samples were analyzed using the HILIC methods described below using 25 mM formate pH 3 (Thermo Scientific, 270480250) for positive mode or 25 mM ammonium carbonate pH 9 (Millipore Sigma, 379999) for negative mode on pump A and distilled 95% Optima LC-MS grade acetonitrile (Fisher Chemical A955-4) on pump B. Positive mode data collection was performed using a SeQuant ZIC-cHILIC 3 μm, 100Å, 100 x 2.1 mm, PEEK-lined stainless-steel column (Millipore Sigma 1.50657) with the following linear gradient conditions: 0-2.50 min, 0.15ml/min, 20% pump A and 80% pump B; 2.50-11.50 min, 0.15ml/min, 20% to 99% pump A and 80% to 1% pump B; 11.50-12.50 min, 0.15ml/min, 99% to 20% pump A and 1% to 80% pump B; 12.50-16.50min, 0.15ml/min, 20% pump A and 80% bump B; 16.50 -21.50 min, 0.45 ml/min, 20% pump A and 80% bump B. The sample injection volume was 2 μL for positive mode. Source conditions for positive mode: ion source gas 1 and 2 40 psi, curtain gas 30 psi, CAD gas 7 psi, source temperature 250°C, spray voltage 5,500 V, declustering potential 50 V, DP speed 0 V, collision energy 10V, and CE spread 0 V. Source conditions for negative mode: ion source gas 1 and 2 40 psi, curtain gas 30 psi, CAD gas 7 psi, source temperature 250°C, spray voltage -4500 V, declustering potential -80 V, DP speed 0 V, collision energy -10 V, and CE spread 0 V. The TOF mass range for positive and negative mode was 50 Da to 1,250 Da with an accumulation time of 0.250 seconds and scan time of 0.278 seconds.The system was routinely calibrated before runs using the SCIEX ESI Positive Calibration Solution for the SCIEX X500B System (SCIEX 5049910) for positive mode and SCIEX ESI Negative Calibration Solution for the SCIEX X500 System (SCIEX 5042913) for negative mode. Instrument sensitivity, mass accuracy, and retention time performance was confirmed before each run by comparative analysis of triplicate injections of parallel-processed, pooled metabolite library to previous standards.

*MIDAS data processing and analysis*
MIDAS LC-MS data were processed using SCIEX OS 3.3 with a targeted method to determine metabolite abundances in protein and metabolite chambers, as previously described (3), with additional normalization and filtering steps detailed below. Raw MS spectra were integrated and extracted ion chromatograms (XICs) were identified based on intact mass, chemical formula, adduct/charge state, precursor mass, and pre-determined retention time. Metabolite abundance was calculated as the integrated area under the XIC peak trace (counts per second). Missing metabolite measurements were explicitly imputed as NA to enable thresholding. Samples with total ion count (TIC) below one-quarter of the median TIC for their LC-MS mode were removed. Ion counts were normalized using a sample-level median polish to correct for global signal differences

across samples. Extremely low or missing values were floored to metabolite-specific limits of detection. For each dialysis replicate, $log_2$-transformed abundances were computed, and $log_2$(fold change) was calculated as the difference between protein and metabolite chambers. Injection replicates were averaged prior to fold-change calculation. Dialysis replicates with missing chamber measurements were excluded. Technical replicates were retained for outlier detection. Metabolites dominated by noise were removed based on comparison of a null model (intercept only) versus a protein-specific model and RMSE > 0.5 across dialysis replicates. Remaining metabolites were classified as "protein signal" and retained for analysis. For each metabolite–protein pair, up to one extreme outlier was removed using a z-score cutoff of 4. Dialysis replicates were then averaged to yield a consensus fold-change per protein–metabolite pair. To remove variation not specific to a given protein–metabolite interaction, the first two principal components of the full screening dataset were projected out on a per-metabolite-pool basis, generating $log_2$(corrected fold change). Protein–metabolite interactions were identified as extreme corrected fold changes relative to each metabolite's distribution across all proteins. For each metabolite, a "no-signal" background distribution was estimated using robust measures of central tendency (median) and variability ($\sigma$ = IQR/1.35). Z-scores were computed for each protein–metabolite pair, and p-values were derived from the standard Normal distribution. False discovery rates were controlled using Storey's q-value method (4). PMIs with $Q < 0.05$ and log2(corrected fold change) > 2 were considered significant.

*Crystallization and X-ray data collection*

FicD crystallization was performed following a protocol with medications (5). $FicD^{E234G}$ (105-433) was expressed and purified as above. Protein was concentrated to ~ 9 mg/mL in the buffer containing 25 mM Tris (pH 8.0), 150 mM NaCl. 5 mM DTT and 5 mM $CaCl_2$ were added before crystallization. Protein crystallization was performed with hanging drop vapor diffusion method. Crystals were obtained at 16 ℃ with a protein:buffer ratio of 2:1 (100 mM Bis-Tris propane (pH 7.5), 200 mM potassium sodium tartrate, and 21% PEG 3350). Soaking of crystals (16°C, overnight) were performed with 10 mM $MgCl_2$ and 1 mM FPP in cryoprotectant buffer (100 mM Bis-Tris propane (pH 7.3), 200 mM potassium sodium tartrate, 21% PEG 3350, and 40% saturated sucrose). Soaked crystals were harvested and frozen in liquid nitrogen. Crystal diffraction data were collected at the Advanced Light Source (ALS) beamline 8.2.2. Data were processed with Xia2 (6) and DIALS (7). Data collection statistics are provided in Table S2.

*Structure refinement*

An AlphaFold structure model of $FicD^{E234G}$ (105-433) was used for molecular replacement in the program Phenix (8). The model was manually adjusted in the program COOT (9). Structure refinement was performed in the program PHENIX (8) . The statistics for refinement of the model are provided in Table S2. Structure figures were prepared with PyMol (The PyMOL Molecular Graphics System, Version 2.4, Schrödinger, LLC.).

*Differential Scanning Fluorimetry*

Thermal shift assays were performed in Biorad CfX384 Touch Real-Time PCR Detection System in Applied Biosystems Microamp Optical 384 well plates (Cat no. 4309849) with MicroAMP Optical Adhesive Film (Cat no. 4313663). Reactions contained 50mM HEPES pH 7.4, 200mM NaCl, 200mM KCl, 200uM DTT, 5X Invitrogen Sypro Orange (Cat no. S6650), 8-9µM protein, and indicated nucleotide, cofactor, or small molecule in a total volume of 20uL. For every experiment 'no protein' controls were also included with the same reaction conditions. All reactions were performed in technical triplicates. Sealed plates containing reaction mixes were centrifuged for 1-5 min at 1000 RPM prior to temperature scanning (FRET channel), which was performed from 6°C to 95°C at a ramp rate of 0.5°C/cycle with 5 second hold per cycle.

DSF data was analyzed with traditional methods automated in a Python package (https://pypi.org/project/instawell/). Briefly, for each temperature, the RFU values of technical triplicates are averaged. For each timepoint, the background fluorescence was subtracted using the RFU values of the 'no protein' control. From these data, the first derivative of the RFU was calculated. The melting temperature ($T_m$) is defined as the first derivative peak value. Data expressed as arbitrary RFU values is min-max scaled to values between 0 and 1 for data visualization. Min-max scaled RFU values and the first derivatives are plotted as a function of temperature in Graphpad Prism 10.6.1. In Prism, the first derivative values are uniformly nudged by -1 to facilitate presenting RFU values and their first derivatives on the same graph.

To generate dose-response curves, $\Delta T_m$, was first calculated by subtracting the $T_m$ of the 'no ligand' control from the $T_m$ of relevant ligand-containing reactions. $\Delta T_m$ was then plotted as a function of ligand concentration. Dose-response curves were fit using GraphPad Prism 10.6.1 with the [Agonist] vs. Response – Variable Slope (four parameters) model. For all dose-response curves, each data point encompassed at least two independent experiments (indicated in figure legends), with each independent experiment containing three technical replicates.

*Isothermal Titration Calorimetry*
Proteins were dialyzed against 25mM HEPES pH 7.4/100mM KCl/10% glycerol prior to titration. Dialyzed proteins were then spun down at 21,000G for 15 minutes at 4C to remove trace insoluble material. The protein concentration was measured spectrophotometrically and then diluted to working concentration with 25mM HEPES pH 7.4/100mM KCl/10% glycerol. $MgCl_2$ was added to proteins to a final concentration of 4mM. ITC experiments were performed in a Micro-Cal PEAQ-ITC (Malvern Panalytical, Worcestershire, UK) calorimeter with a stirred 206.2 µL reaction cell held at 20 °C. 250µM FPP was titrated into 25uM $\Delta$104hFic$^{L258D}$ with 750rpm stirring. The first injection was 0.5 µL, followed by twenty 1.9 µL injections. ITC data were integrated and baseline corrected using NITPIC 2.1.5 (10, 11). Integrated data was globally analyzed in SEDPHAT 15.b (12) using a model considering a single class of binding sites. Thermogram and binding figures were plotted in GUSSI 2.1.6 (13).

*In-vitro activity assays*
For AMPylation assays with FicD and BiP, $\Delta$104FicD$^{WT}$ and $\Delta$26hBiP$^{T229A}$ were combined at a ratio of 1:80 in buffer containing 25mM HEPES pH 7.4, 100mM KCl, 5mM MgCl2, 500µM ATP, 1mM DTT, and small molecule where indicated. Assays with $\Delta$104FicD$^{WT}$ were incubated for 1 hour. $\Delta$104FicD$^{R371S}$ and $\Delta$104FicD$^{R374H}$ have very weak AMPylation activity and therefore, contained 10X the amount of FicD as in reactions containing FicD$^{WT}$ and were incubated at 37C for 4.5 hours. Reactions were stopped with the addition of EDTA and 5X SDS sample buffer (250mM Tris pH 6.8, 10% Lithium dodecyl sulfate, 50% glycerol, 50mM DTT, 0.05% Bromophenol blue) and then heated at 98°C for 5 minutes.

For deAMPylation assays with FicD and BiP, $\Delta$26hBiP$^{T229A}$ was in-vitro AMPylated by $\Delta$104hFicD$^{E234G/L258D}$ at a ratio of 1:320 in buffer containing 25mM HEPES pH 7.4, 100mM KCl, 5mM MgCl2, 500uM ATP, 0.5% CHAPS, 1mM DTT. After one hour of incubation at 30-37C, the entire reaction mix was transferred into an Amicon 0.5mL 50kDa MWCO concentrator (cat no. UFC5010) and spun down at 14,000 x g at 4C for 5-10 minutes. 400uL-500µL of 50mM Tris pH 7.4/150mM NaCl/10% glycerol/1mM DTT was added to the concentrated AMPylation reaction and then spun down again at 14,000 x g at 4C for 5-10 minutes. This step was repeated four times to buffer exchange AMPylated $\Delta$26hBiP$^{T229A}$.

$\Delta$104hFicD$^{WT}$ and $\Delta$26hBiP$^{T229A\text{-}AMP}$ were combined at a ratio of 1:80 in buffer containing 25mM HEPES pH 7.4, 100mM KCl, 5mM MgCl2, 1mM DTT and the indicated small molecule where applicable. Reactions were incubated at 37C for 1 hour and stopped by the addition EDTA. 5X SDS sample buffer (250mM Tris pH 6.8, 10% Lithium dodecyl sulfate, 50% glycerol, 50mM DTT, 0.05% Bromophenol blue) was then added to the reactions and reactions were heated for 5 minutes at 95°C.

For in-vitro AMPylation assays with VopS and CDC42, $\Delta$30VopS and GST-CDC42 – provided by a previous lab member – were combined at a 1:80 molar ratio in 50mM Tris pH 7.4/100mM NaCl/1mM DTT in the presence of GPP or FPP, where applicable. Reactions were started by the addition of ATP/MgCl2 at a final concentration of 500µM/5mM and reactions were incubated for 1 hour at room temperature. Reactions were stopped by the addition of 5X SDS sample buffer

(250mM Tris pH 6.8, 10% Sodium dodecyl sulfate, 50% glycerol, 50mM DTT, 0.05% Bromophenol blue) and heated at 98°C for 5 minutes.

Rhamnosylation assays were performed as previously described (2). Briefly, AvrB and GST-Rin4 were combined in the presence of GPP or FPP, where applicable. Reactions were started by the addition of UDP-Rhamnose at a final concentration of 200M and reactions were incubated for 1 hour at room temperature. Reactions were stopped by the addition of 5X SDS sample buffer (250mM Tris pH 6.8, 10% Sodium dodecyl sulfate, 50% glycerol, 50mM DTT, 0.05% Bromophenol blue) and heated at 98°C for 5 minutes.

*Immunoblotting*
Equal volumes of samples suspended in 5X SDS sample buffer were resolved on hand-cast 10% 1.5mm Tris-Glycine gels with the BioRad Mini-PROTEAN Tetra Cell system. Proteins were transferred onto Millipore Immobilon-P 0.45µM Pore size PVDF membrane (cat no. IPVH00010) for 10 min at 1.5 Amps with Invitrogen Powerblotter Semi-Dry transfer system. Membranes were then dried at 37C for 10-15 minutes immediately after transfer. Membranes were rewetted with 100% methanol and rocked in appropriate blocking buffer (5% nonfat dry milk or 5% RPI Bovine Serum Albumin Fraction V (cat no. A30075-100.0) in 1X TTBS for 1 hour at room temperature. Primary antibodies were diluted in appropriate blocking buffer and then rocked with membranes 1 hour at room temperature. Membranes were then washed in 1X TTBS three to four times. Secondary antibodies were diluted in appropriate blocking buffer and then rocked with membranes overnight at 4C. Membranes were then washed in 1X TTBS three to four times and developed with Advansta WesternBright ECL Spray (cat no. K-12049-D50) or ThermoScientific SuperSignal West Femto Maximum Sensitivity Substrate (cat no. 34096). Membranes were then imaged in the BioRad ChemiDoc Imaging System.

*Intact Mass*
Intact mass analysis of proteins was performed by the UT Southwestern Proteomics Core. Intact protein samples were analyzed by LC/MS, using a Sciex X500B QTOF mass spectrometer operating in positive ion mode, coupled to an Agilent 1290 Infinity II HPLC. Samples were injected onto a POROS R1 reverse-phase column (2.1 x 30 mm, 20 µm particle size, 4000 Å pore size) and desalted. The mobile phase flow rate was 300 µL/min and the gradient was as follows: 0-3 min: 0% B, 3-4 min: 0-15% B, 4-16 min: 15-55% B, 16-16.1 min: 55-80% B, 16.1-18 min: 80% B. The column was then re-equilibrated at initial conditions prior to the subsequent injection. Buffer A contained 0.1% formic acid in water and buffer B contained 0.1% formic acid in acetonitrile.

The mass spectrometer was controlled by Sciex OS v.3.0 using the following settings: Ion source gas 1 30 psi, ion source gas 2 30 psi, curtain gas 35, CAD gas 7, temperature 300 oC, spray voltage 5500 V, declustering potential 125 V, collision energy 10 V. Data was acquired from 400-2000 Da with a 0.5 s accumulation time and 4 time bins summed. The acquired mass spectra for the proteins of interest were deconvoluted using BioPharmaView v. 3.0.1 in order to obtain the molecular weights. Peaks were deconvoluted from 39-43 kDa, with 10 iterations, signal to noise threshold >5, resolution = 2500.

**SI References**


1. D. Höpfner *et al.*, Monoclonal Anti-AMP Antibodies Are Sensitive and Valuable Tools for Detecting Patterns of AMPylation. *iScience* **23**, 101800 (2020).
2. W. Peng *et al.*, Pseudomonas effector AvrB is a glycosyltransferase that rhamnosylates plant guardee protein RIN4. *Sci Adv* **10**, eadd5108 (2024).
3. K. G. Hicks *et al.*, Protein-metabolite interactomics of carbohydrate metabolism reveal regulation of lactate dehydrogenase. *Science* **379**, 996-1003 (2023).
4. J. D. Storey, R. Tibshirani, Statistical significance for genomewide studies. *Proc Natl Acad Sci U S A* **100**, 9440-9445 (2003).

5. T. D. Bunney *et al.*, Crystal structure of the human, FIC-domain containing protein HYPE and implications for its functions. *Structure* **22**, 1831-1843 (2014).
6. G. Winter, xia2: an expert system for macromolecular crystallography data reduction. *J Appl Crystallogr* **43**, 186-190 (2010).
7. G. Winter *et al.*, DIALS: implementation and evaluation of a new integration package. *Acta Crystallogr D Struct Biol* **74**, 85-97 (2018).
8. P. D. Adams *et al.*, PHENIX: a comprehensive Python-based system for macromolecular structure solution. *Acta crystallographica. Section D, Biological crystallography* **66**, 213-221 (2010).
9. P. Emsley, B. Lohkamp, W. G. Scott, K. Cowtan, Features and development of Coot. *Acta crystallographica. Section D, Biological crystallography* **66**, 486-501 (2010).
10. S. Keller *et al.*, High-precision isothermal titration calorimetry with automated peak-shape analysis. *Anal Chem* **84**, 5066-5073 (2012).
11. T. H. Scheuermann, C. A. Brautigam, High-precision, automated integration of multiple isothermal titration calorimetric thermograms: new features of NITPIC. *Methods* **76**, 87-98 (2015).
12. C. A. Brautigam, H. Zhao, C. Vargas, S. Keller, P. Schuck, Integration and global analysis of isothermal titration calorimetry data for studying macromolecular interactions. *Nat Protoc* **11**, 882-894 (2016).
13. C. A. Brautigam, Calculations and Publication-Quality Illustrations for Analytical Ultracentrifugation Data. *Methods Enzymol* **562**, 109-133 (2015).

### Figures

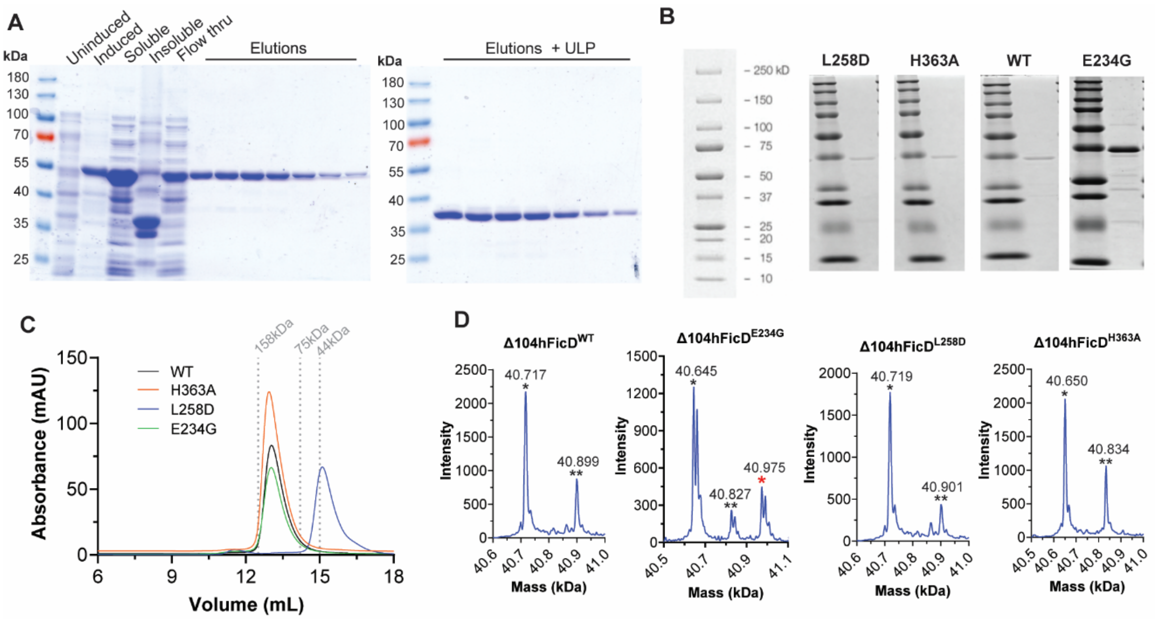


**Fig S1. Purification of Δ104hFicD. (A)** SDS-PAGE of samples from representative Δ104hFicD$^{WT}$ purification. Human FicD$^{WT}$ is expressed as an N-terminal 6xHis-Sumo fusion. Bacterial samples are taken before (uninduced) and after IPTG induction (induced). Bacterial lysate is centrifuged to separate soluble material (soluble) from insoluble material (insoluble). Δ104hFicD is purified by NiNTA affinity from the soluble lysate and eluted with imidazole (elutions). IPTG induction produces excess FicD, which is seen in the lysate post-IMAC (flow-thru). Elutions are then cut with ULP protease (Elution + ULP) prior to cleanup by gel filtration. Results are representative of all FicD mutants. **(B)** SDS-PAGE of Δ104hFicD proteins screened by MIDAS. **(C)** FPLC chromatogram of FicD mutants. 600ug-800ug of Δ104hFicD$^{WT}$ (black), Δ104hFicD$^{E234G}$ (green), Δ104hFicD$^{L258D}$ (blue), and Δ104hFicD$^{H363A}$ (orange) were gel filtered with 50mM Tris pH 8/150mM NaCl in a Cytiva Superdex-200 increase 10/300 GL column. Dotted lines indicate elution volumes of gel filtration standards Ovalbumin (44kDa), Conalbumin (75kDa), and Aldolase (158kDa). **(D)** Intact mass of proteins screened by MIDAS. * indicates mass peak close to the theoretical mass (WT – 40.717kDa, E234G – 40.645, L258D – 40.719kDa, H363A – 40.651kDa) ** indicates +183 mass shift likely associated with AEBSF from protease inhibitor cocktail used in IMAC purification. Asterisks * in red indicates +329 mass shift associated with AMPylation

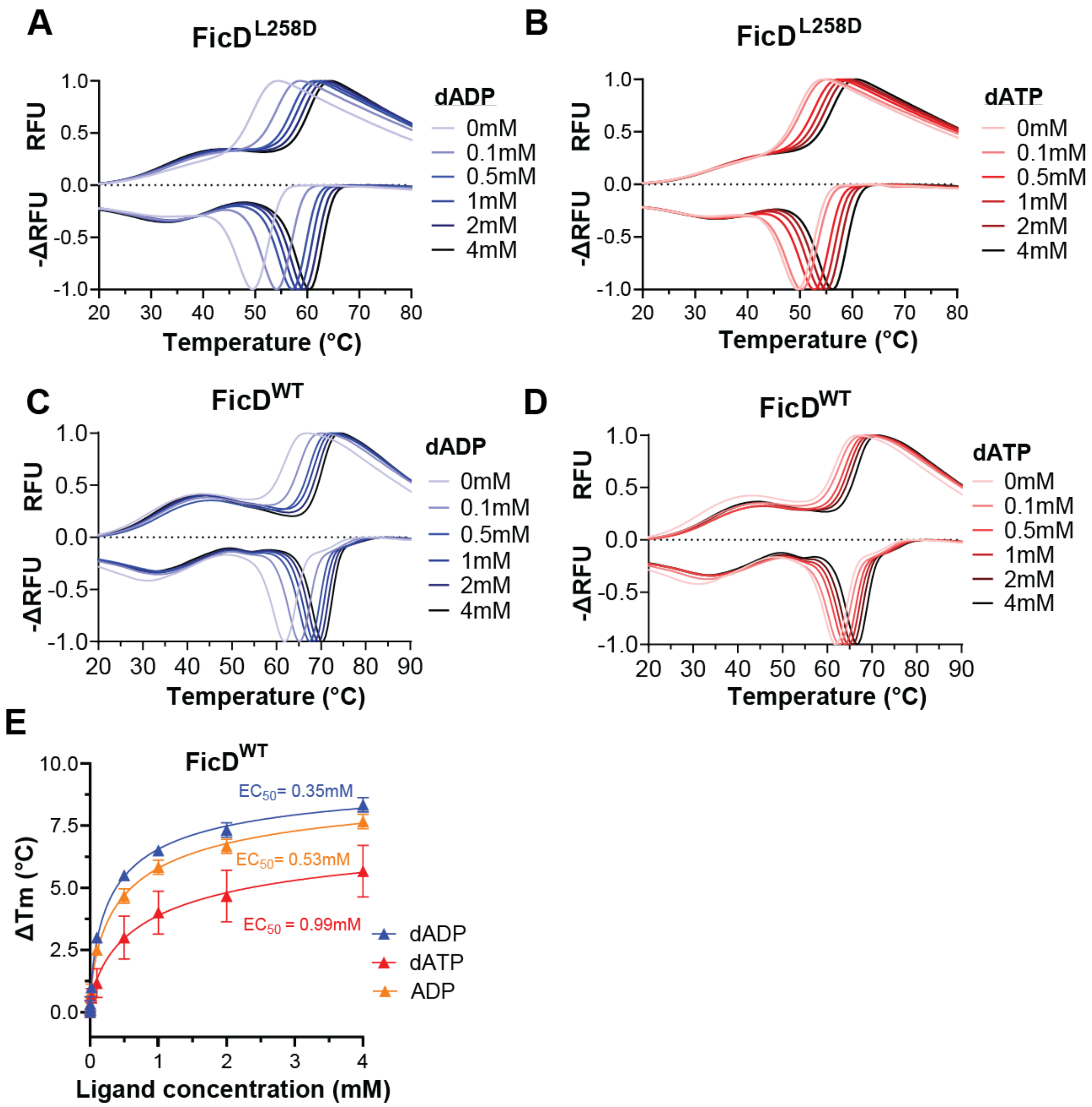


**Fig S2. FicD interaction with analogues of canonical ligands (A-D)** Representative melt curves of FicD measured in relative fluorescence units (RFU, upper) and the derivative (-ΔRFU, lower) in the presence of increasing concentrations of nucleotides (colorscaled from light to dark) for (**A**) FicD$^{L258D}$ and dADP, **(B)** FicD$^{L258D}$ and dATP, **(C)** FicD$^{WT}$ and dADP, or **(D)** FicD$^{WT}$ and dATP. **(E)** Nucleotide-induced DSF melting point analysis of FicD$^{WT}$ in response to dADP (blue), dATP (red), and ADP (orange). Each data point encompasses three independent experiments. EC50 values were calculated in GraphPad Prism 10.6.1 using the [Agonist] vs. Response – Variable Slope (four parameters) model.

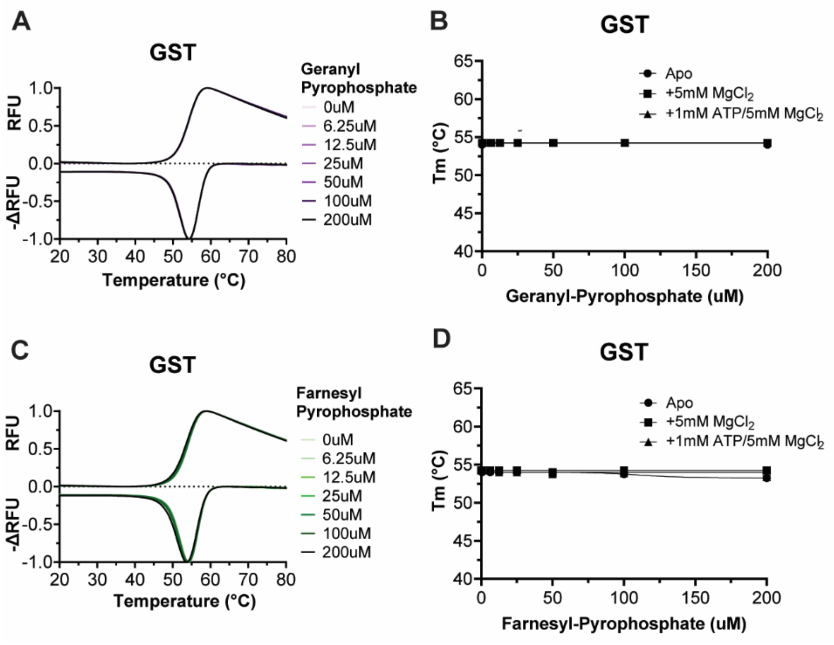


**Fig S3. DSF with GST and GPP/FPP. (A)** Representative melt curves of GST measured in (RFU, upper) and the derivative (ΔRFU, lower) at the indicated concentrations of GPP. All curves are plotted, but not visible due to overlap **(B)** GPP-induced DSF melting point analysis of GST. Each data point encompasses two independent experiments. All data is plotted but may not visible due to overlap. All error bars are plotted but may be occluded by the data point symbol or not present if there was no error between experiments. **(C)** Representative melt curves of GST measured in (RFU, upper) and the derivative (ΔRFU, lower) at the indicated concentrations of FPP (RFU = relative fluorescence units). **(D)** FPP-induced DSF melting point analysis of GST in response to FPP. Each data point encompasses two independent experiments.

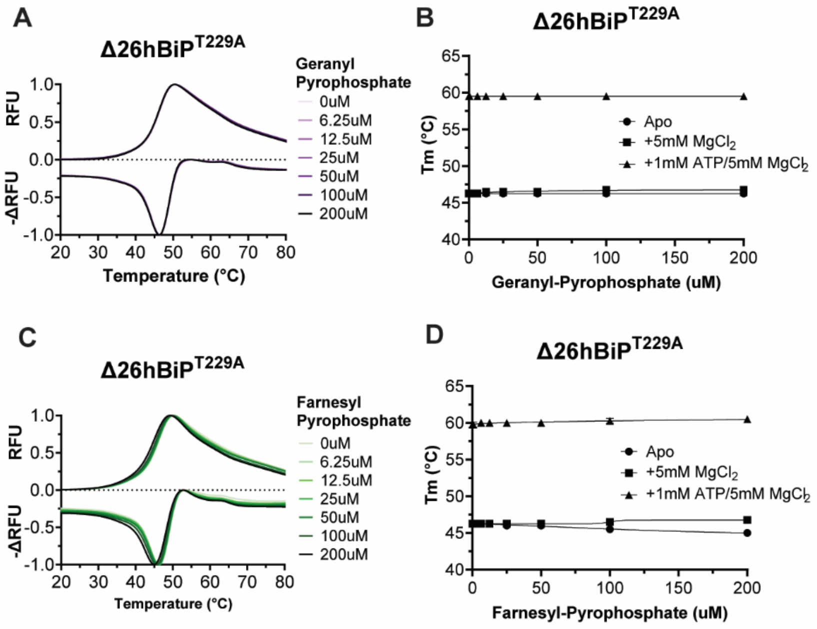


**Fig S4. DSF with BiP and GPP/FPP. (A)** Representative melt curves of Δ26hBiP$^{T229A}$ measured in (RFU, upper) and the derivative (ΔRFU, lower) at the indicated concentrations of GPP **(B)** GPP-induced DSF melting point analysis of Δ26hBiP$^{T229A}$(Apo). Each data point encompasses two independent experiments. All error bars are plotted but may be occluded by the data point symbol or not present if there was no error between experiments. **(C)** Representative melt curves of Δ26hBiP$^{T229A}$ (Apo) measured in (RFU, upper) and the derivative (ΔRFU, lower) at the indicated concentrations of FPP. **(D)** FPP-induced DSF melting point analysis of Δ26hBiP$^{T229A}$. Each data point encompasses two independent experiments.

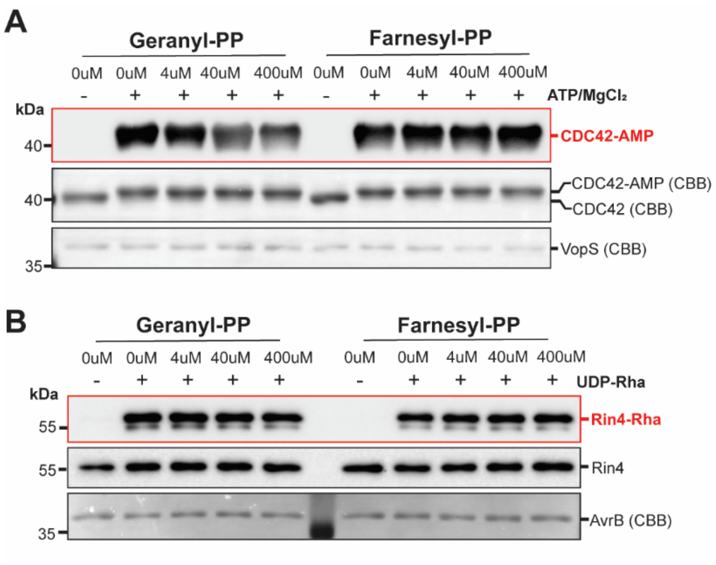


**Fig S5. Assessment of GPP/FPP effects on bacterial Fido proteins. (A)** In-vitro AMPylation assay with VopS and CDC42. GST-CDC42 was AMPylated by Δ30VopS in the presence of the indicated concentration of GPP or FPP. GST-CDC42$^{AMP}$ was visualized with immunodetection and the shift in mobility visible by Coomassie brilliant blue stain (CBB). VopS was visualized by Coomassie brilliant blue stain (CBB) **(B)** In-vitro Rhamnosylation assay with AvrB and Rin4. GST-Rin4 was Rhamnosylated by AvrB in the presence of the indicated concentration of GPP or FPP. Rin4$^{T166\text{-}Rha}$ and total Rin4 were visualized with immunodetection and AvrB was quantified by Coomassie Brilliant Blue staining (CBB).

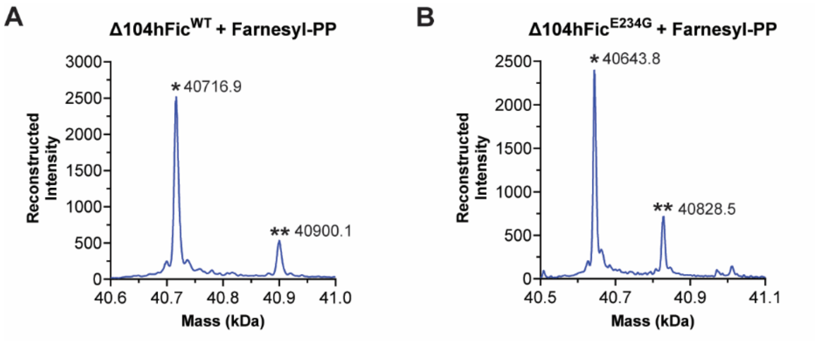


**Fig S6. Total mass of FicD incubated with Farnesyl Pyrophosphate. (A)** $FicD^{WT}$ was incubated with 50uM Farnesyl-Pyrophosphate and 5mM $MgCl_2$. *indicates mass peaks close to the theoretical mass (40.717 kDa for $FicD^{WT}$) ** indicates +183 mass shift likely associated with AEBSF from protease inhibitor cocktail used in IMAC purification. **(B)** $FicD^{E234G}$ was 50uM Farnesyl-Pyrophosphate and 5mM $MgCl_2$. * in black indicates mass peaks close to the theoretical mass (40.645kDa for $\Delta 104hFicD^{E234G}$) ** indicates +183 mass shift likely associated with AEBSF from protease inhibitor cocktail used in IMAC purification.

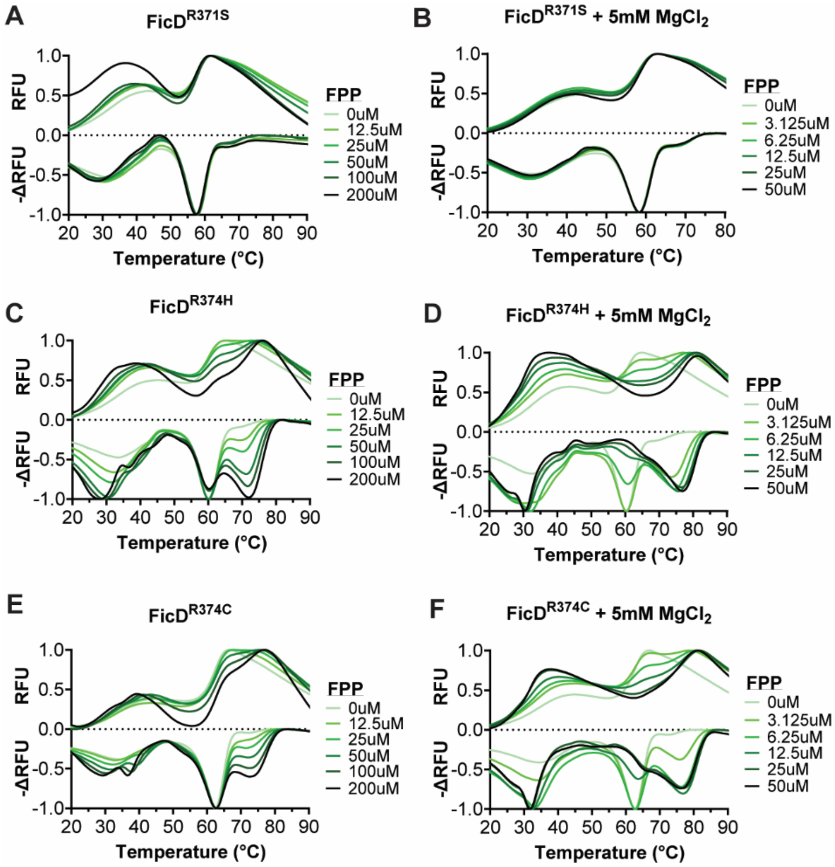


**Fig S7**. **FPP binding to pathogenic FicD variants.** Representative melt curves measured in relative fluorescence units (RFU, upper) and the derivative (-ΔRFU, lower) for FicD variants in the presence of increasing FPP concentrations: **(A)** FicD$^{R371S}$ **(B)** FicD$^{R371S}$ with 5mM MgCl2 **(C)** FicD$^{R374H}$ **(D)** FicD$^{R374H}$ with 5mM MgCl2 and **(E)** FicD$^{R374C}$ and **(F)** FicD$^{R374C}$ with 5mM MgCl2.

### Tables

#### Table S1. Primers used to clone hFicD and hBiP

| Construct/Mutant | Forward Primer Sequence | Reverse Primer Sequence |
|---|---|---|
| Δ104hFic_BamHI_NotI | 5'-CGCGGATCCGAAGCCAGAGCTGCCCTGAACC-3' | 5'-GAGTGCGGCCGCTTAGGGCTTCACAGGAAGCGTC-3' |
| Δ26hBiP_BamHI_NotI | 5'-CGCGGATCCGTGGGCACGGTGGTCGGC-3' | 5'-GAGTGCGGCCGCCTACAACTCATCTTTTTCTGCTGTATCCTCTTC-3' |
| hFic_E234G | 5'-GGGGGCAACACCCTCACCCTCTCGG-3' | 5'-GATGGCCACCGTGTGGTAGATGTGG-3' |
| hFic_L258D | 5'-GATGAGGAGCAGAACGAGGTCATAGG-3' | 5'-GCTCTTCCCGGGCACGGCGTAGCGGG-3' |
| hFic_H363A | 5'-GCCCCTTTCATTGATGGCAACGGGAG-3' | 5'-GATGTAAACGAGTTTATAATGGGC-3' |
| hFic_R371S | 5'-AGCACCTCCCGTCTGCTCATG-3' | 5'-CCCGTTGCCATCAATGAAAGGGTG-3' |
| hFic_R374H | 5'-CATCTGCTCATGAACCTCATCCTCATG-3' | 5'-GGAGGTCCTCCCGTTGCCATC-3' |
| hFic_R374C | 5'-TGTCTGCTCATGAACCTCATCCTC-3' | 5'-GGAGGTCCTCCCGTTGCCATC-3' |
| hBiP_T229A | 5'-GCCTTCGATGTGTCTCTTCTCACCATTG-3' | 5'-TCCGCCACCCAGGTCAAACACCAGG-3' |
| 105-433hFic | 5'-TAAGCGGCCGCACTCGAGCAC-3' | 5'-TGTGGCAAAAAGCAGGGTGTCCAG-3' |

**Table S2. MIDAS hits for Δ104hFicD$^{WT}$**

| Mass_RT_ID | Metabolite Name | Log2(corrected_fold_change) | q-Value |
|---|---|---|---|
| 685 | Deoxyadenosine triphosphate | 4.489066429 | 0.001661423 |
| 621 | Adenosine 3',5'-diphosphate OR dGDP OR ADP | 3.343909179 | 3.66E-04 |
| 796 | 3,3'-Diiodothyronine | 3.242622698 | 7.71E-04 |
| 846 | Farnesyl pyrophosphate | 3.222110512 | 0.001860495 |
| 680 | dADP | 3.081073144 | 3.45E-30 |
| 865 | 3,3'-Diiodothyronine | 3.080746404 | 0.002099897 |
| 622 | Farnesyl pyrophosphate | 2.971253285 | 5.49E-12 |
| 737 | 3,3'-Diiodothyronine | 2.939650049 | 0.00780651 |
| 659 | Geranyl-PP | 2.844244414 | 5.70E-56 |
| 810 | Phosphorylcholine | 2.726798047 | 2.51E-08 |
| 585 | Lithocholic acid glycine conjugate | 2.555224294 | 0.038943658 |
| 598 | Lithocholic acid | 2.551945791 | 0.014578573 |
| 400 | L-alpha-Aminobutyric acid OR gamma-Aminobutyric acid OR 3-Aminoisobutanoic acid OR Isoleucyl-Glycine | -3.175600673 | 4.83E-172 |
| 467 | Glyoxylic acid | -3.454940275 | 3.32E-125 |

**Table S3. MIDAS hits for Δ104hFicD$^{E234G}$**

| Mass_RT_ID | Metabolite Name | Log2(corrected_fold_change) | q-Value |
|---:|---|---:|---:|
| 71 | Acetylcholine | 5.844361025 | 1.61E-205 |
| 796 | 3,3'-Diiodothyronine | 2.644186312 | 0.011788269 |
| 291 | 3,3'-Diiodothyronine OR 3,5-Diiodothyronine | 3.093127643 | 0.002359772 |
| 865 | 3,3'-Diiodothyronine | 2.487148443 | 0.023745028 |
| 180 | 3,3'-Diiodothyronine OR 3,5-Diiodothyronine | 2.746801516 | 0.008839425 |
| 737 | 3,3'-Diiodothyronine | 2.494570871 | 0.037103428 |
| 877 | 3'-Monoiodo-L-thyronine | 2.322543023 | 0.004675749 |
| 680 | dADP | 2.859882938 | 4.62E-26 |
| 388 | Uridine 5'-monophosphate OR+A2:D20 Uridine 3'-monophosphate | 2.079763504 | 1.97E-167 |
| 882 | Adenosine 3',5'-diphosphate OR dGDP OR ADP | 2.634634946 | 3.56E-04 |
| 621 | Adenosine 3',5'-diphosphate OR dGDP OR ADP | 2.648318931 | 1.10E-02 |
| 855 | Flavin mononucleotide | 2.084562916 | 5.87E-03 |
| 852 | Geranyl-PP | 2.314812875 | 9.57E-05 |
| 716 | IDP | 2.529694912 | 0.00000451 |
| 685 | Deoxyadenosine triphosphate | 3.342502867 | 0.04941929 |
| 587 | Mandelic acid OR 3,4-Dihydroxyphenylacetaldehyde OR p-Hydroxyphenylacetic acid | -3.54781522 | 1.97E-283 |
| 634 | Isovaleryl-CoA | -3.150135628 | 4.44E-08 |
| 784 | Isovaleryl-CoA | -3.290864991 | 2.02E-06 |
| 268 | Uridine 5'-monophosphate OR Uridine 3'-monophosphate | -7.356036567 | 0.00E+00 |

**Table S4. MIDAS hits for Δ104hFicD$^{L258D}$**

| Mass_RT_ID | Metabolite Name | Log2(corrected_fold_change) | q-Value |
|---|---|---|---|
| 810 | Phosphorylcholine | 4.026657953 | 1.80E-17 |
| 622 | Farnesyl pyrophosphate | 3.430081372 | 6.03E-16 |
| 846 | Farnesyl pyrophosphate | 3.270500539 | 0.001494892 |
| 659 | Geranyl-PP | 3.248792222 | 2.76E-73 |
| 716 | IDP | 3.173357999 | 1.28E-09 |
| 796 | 3,3'-Diiodothyronine | 3.097224609 | 0.001581634 |
| 865 | 3,3'-Diiodothyronine | 2.657183967 | 0.012552777 |
| 737 | 3,3'-Diiodothyronine | 2.614952966 | 0.024976034 |
| 680 | dADP | 2.437749264 | 4.94E-19 |
| 625 | Adenosine phosphosulfate | 2.029900916 | 0.004440704 |
| 691 | Methylsuccinic acid | -3.633930028 | 6.07E-21 |
| 35 | D-Arabitol OR Ribitol | -5.940490662 | 0.00E+00 |

**Table S5. MIDAS hits for Δ104hFicD$^{H363A}$**

| Mass_RT_ID | Metabolite Name | Log2(corrected_fold_change) | q-Value |
|---|---|---|---|
| 846 | Farnesyl pyrophosphate | 5.139122794 | 1.69E-08 |
| 622 | Farnesyl pyrophosphate | 3.152695213 | 1.75E-13 |
| 796 | 3,3'-Diiodothyronine | 2.431780899 | 0.026726808 |
| 621 | Adenosine 3',5'-diphosphate OR dGDP OR ADP | 2.290181332 | 0.044795438 |
| 659 | Geranyl-PP | 2.219796903 | 7.85E-34 |
| 400 | L-alpha-Aminobutyric acid OR gamma-Aminobutyric acid OR 3-Aminoisobutanoic acid OR Isoleucyl-Glycine | 2.045593094 | 8.65E-75 |
| 852 | Geranyl-PP | 1.669026472 | 0.014261334 |
| 691 | Methylsuccinic acid | -2.752610589 | 2.34E-12 |

**Table S6 | Data collection and refinement statistics.**

| **Data collection** | |
|---|---|
| Crystal | FicD$^{E234G}$-FPP |
| Wavelength (Å) | 1.0000 |
| Resolution range (Å) | 48.94 - 2.58 (2.64 - 2.58) |
| Space group | $P12_11$ |
| Unit cell | 71.02, 76.98, 91.33 90, 107.2, 90 |
| Unique reflections | 29,695 (2,114) |
| Completeness (%) | 99.59 (98.83) |
| Mean I/sigma(I) | 7.3 (0.8) |
| Wilson B-factor (Å$^2$) | 7.21 |
| R-merge (%) | 4.6 |
| $CC_{1/2}$ | 0.990 (0.299) |
| **Refinement** | |
| Reflections for refinement | 29,695 (2,114) |
| Reflections for R-free | 1,999 (143) |
| R-work (%) | 23.33 (39.76) |
| R-free (%) | 29.05 (45.58) |
| Atoms (non-hydrogen) | 5,462 |
| macromolecules | 5,328 |
| ligands | 50 |
| solvent | 84 |
| R.m.s.d. bond (Å) | 0.010 |
| R.m.s.d. angle (°) | 1.13 |
| Ramachandran favored (%) | 94.95 |
| Ramachandran allowed (%) | 4.74 |
| Ramachandran outliers (%) | 0.31 |
| Rotamer outliers (%) | 4.67 |
| Clashscore | 16.90 |
| Average B-factor (Å$^2$) | 61.02 |
| macromolecules | 61.33 |
| ligands | 44.85 |
| solvent | 50.78 |

Statistics for the highest-resolution shell are shown in parentheses. $R_{merge}=\Sigma_h\Sigma_i|I_{h,i}-I_h|/\Sigma_h\Sigma_i I_{h,i}$, where $I_h$ is the mean intensity of the $i$ observations of symmetry related reflections of $h$. $R=\Sigma|F_{obs}-F_{calc}|/\Sigma F_{obs}$, where $F_{calc}$ is the calculated protein structure factor from the atomic model).

**Dataset S1 (separate file).** Complete MIDAS screening dataset for Δ104hFicD$^{WT}$.

**Dataset S1 (separate file).** Complete MIDAS screening dataset for Δ104hFicD$^{E234G}$

**Dataset S1 (separate file).** Complete MIDAS screening dataset for Δ104hFicD$^{L258D}$

**Dataset S1 (separate file).** Complete MIDAS screening dataset for Δ104hFicD$^{H363A}$
